# Comparative analysis of the CDR loops of antigen receptors

**DOI:** 10.1101/709840

**Authors:** Wing Ki Wong, Jinwoo Leem, Charlotte M. Deane

**Affiliations:** Department of Statistics, University of Oxford, 24-29 St Giles, Oxford OX1 3LB

## Abstract

The adaptive immune system uses two main types of antigen receptors: T-cell receptors (TCRs) and antibodies. While both proteins share a globally similar *β*-sandwich architecture, TCRs are specialised to recognise peptide antigens in the binding groove of the major histocompatibility complex, while antibodies can bind an almost infinite range of molecules. For both proteins, the main determinants of target recognition are the complementarity-determining region (CDR) loops. Five of the six CDRs adopt a limited number of backbone conformations, known as the ‘canonical classes’; the remaining CDR (*β*3 in TCRs and H3 in antibodies) is more structurally diverse. In this paper, we first update the definition of canonical forms in TCRs, build an auto-updating sequence-based prediction tool (available at http://opig.stats.ox.ac.uk/resources) and demonstrate its application on large scale sequencing studies. Given the global similarity of TCRs and antibodies, we then examine the structural similarity of their CDRs. We find that TCR and antibody CDRs tend to have different length distributions, and where they have similar lengths, they mostly occupy distinct structural spaces. In the rare cases where we found structural similarity, the underlying sequence patterns for the TCR and antibody version are different. Finally, where multiple structures have been solved for the same CDR sequence, the structural variability in TCR loops is higher than that in antibodies, suggesting TCR CDRs are more flexible. These structural differences between TCR and antibody CDRs may be important to their different biological functions.

## 1 Introduction

The adaptive immune system defends the host organism against a wide range of foreign molecules, or antigens, using two types of receptors: T-cell receptors (TCRs) and antibodies (23). TCRs typically recognise peptide antigens presented via the major histocompatibility complex (MHC; 44), while antibodies can bind almost any antigen, including proteins, peptides, and haptens (46). Despite their different roles in the immune response, these proteins share a *β*-sandwich fold (Figure 1; 16, 51).

**Figure 1:**
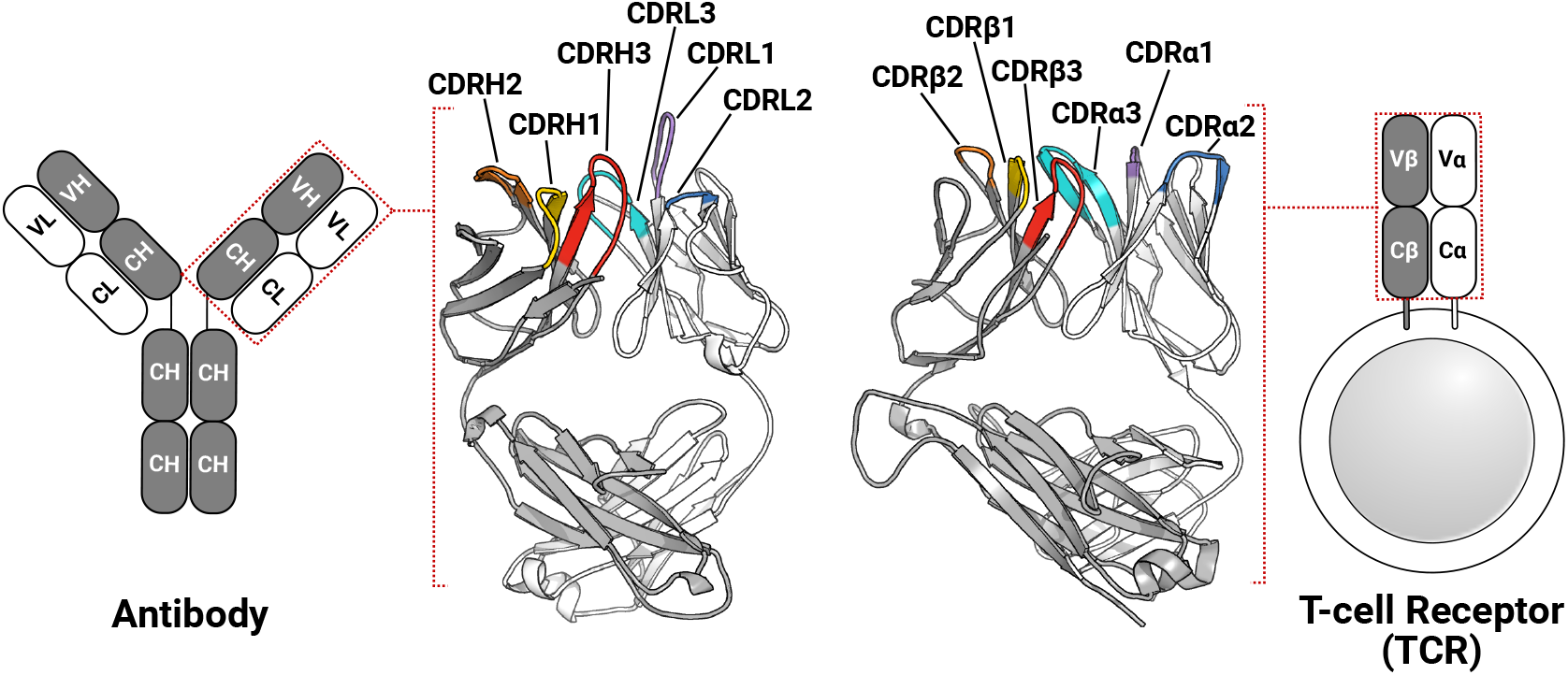
TCRs and antibodies share a globally similar structure. Both proteins are heterodimers, characterised by a set of six CDRs that form the majority of the binding site. Comparable chains and CDRs share colouring schemes; for example, the TCR*β* and antibody heavy chain are coloured grey, while CDR*α*1 and CDRL1 are coloured purple.

In humans, most TCRs are *αβ*TCRs, consisting of one TCR*α* chain and one TCR*β* chain, while most antibodies are comprised of two heavy(H)-light(L) chain dimers (Figure 1). All four types of chains (*α*,*β*, H and L) are formed from the somatic rearrangement of the respective *V*,*D*, and *J* genes of the TCR or antibody loci. The random combination of these genes, alongside further diversification mechanisms (*e.g.* random nucleotide addition), are estimated to yield trillions of unique TCRs and antibodies (5, 20). TCR*α* and antibody light chains are made from the *V* and *J* genes, while TCR*β* and antibody heavy chains are assembled from the *V*,*D*, and *J* genes, making the L-chain equivalent to *α*-chain and H-chain equivalent to *β*-chain (5, 45). In both types of antigen receptors, sequence and structural diversity is concentrated in six hypervariable loops, known as the complementarity determining regions (CDRs). There are three in the TCR*α* chain (CDR*α*1–CDR*α*3) and three in the TCR*β* chain (CDR*β*1–CDR*β*3). Likewise, the light chain and heavy chain of antibodies have three CDRs each (CDRL1–CDRL3, CDRH1–CDRH3). In TCRs, CDR1 and CDR2 typically contact the MHC’s conserved *α*-helices (8, 48), while the CDR3 almost always contacts the peptide antigen (11, 21). All six antibody CDRs can be involved in antigen recognition (26, 46), though the CDRH3 loop is often the most important (34, 53).

A standard way to describe the structures of CDRs is the ‘canonical class’ model, which was first proposed for antibodies in 1987 (9), and has been revisited several times since (e.g. 9, 28, 36, 39, 40). The canonical class model is based on the observation that CDRs adopt a limited number of backbone conformations, and that these conformations can be predicted using sequence features, such as the presence of specific amino acids within or near the CDR loop (e.g. 36, 39, 40). Canonical forms have been used for *in silico* antibody design (29, 41), and predicting the structures of CDR sequences from next-generation sequencing (NGS) datasets (25, 40). Despite the value of canonical classes to antibody design and development, only two studies have so far applied the concept to TCR CDRs (2, 24).

The first clustering of TCR CDR loops was carried out using only seven TCR structures (2). At that time, four canonical classes were identified for CDR*α*1, four for CDR*α*2, three for CDR*β*1, and three classes for CDR*β*2; neither CDR3 loop was clustered. More recently, Klausen *et al.* clustered the CDRs from a non-redundant set of 105 *αβ*TCR and 11 unpaired TCR structures. They performed the clustering in torsion space using an affinity propagation algorithm. In total, 38 canonical forms were characterised. These clusters were then used to construct a sequence-based, random forest classifier, with canonical form prediction accuracies between 63.21% and 98.25% (24).

CDR*β*3 in TCRs and CDRH3 in antibodies show higher variability in sequence composition and structure than the other CDRs (23). While no canonical forms have been defined for CDRH3, several groups have analysed the kinked and extended (or bulged and non-bulged) conformations at the start and end of the loop, known as the ‘base’ or ‘torso’ region (19, 27, 38, 39, 49, 50, 54). Weitzner et al. (54) showed that pseudo bond angle *τ* and pseudo dihedral angle *α* of the second last residue of the CDRH3 loop (Chothia (1) position 101, IMGT (31) position 116) can differentiate between the extended and kinked torso conformations. Finn et al. (19) analysed the first three and last four residues of CDRH3 loops and observed that for the same IMGT position 116, the *ϕ*/*ψ* angles are different in kinked and extended torsos. In this paper, we carry out the first examination of the conformation of the base region of CDR*β*3 loops.

Although TCRs and antibodies are derived from similar genetic mechanisms and share a similar architecture, only a handful of studies have compared them (*e.g.*3, 6, 8, 16, 31, 42). Furthermore, analyses have largely focussed on sequence-based features. For instance, Rock *et al.* found that the CDR*α*3 and CDR*β*3 loops have a different length distribution to CDRL3 and CDRH3 (42), while Blevins *et al.* observed that the TCR CDR1 and CDR2 sequences have more charged amino acids than the analogous antibody CDR1 and CDR2 (8). Given the similarity in the genetic mechanisms, fold and the limited conformational variability in the canonical CDRs, it might be reasonable to expect structural similarity between TCRs and antibodies. Comparing TCRs and antibodies should identify potential characteristics that may inspire antibody-like TCR design, TCR-mimic antibodies and soluble TCRs (14, 16, 52, 57). Such analyses can also highlight structural signatures that may relate to their different biological functions, such as MHC restriction in TCRs (5) and the virtually unconstrained antigen binding in antibodies (46).

In this manuscript, we first used a length-independent clustering method to update the canonical classes of TCR CDRs (40). We then built a sequence-based TCR CDR canonical form prediction algorithm based on an adapted position-specific scoring matrix (55). Next, we attempted to enrich the TCR CDR dataset with antibody CDR structures. We found that TCR and antibody CDRs occupy distinct areas of structural space. In the small number of common conformational clusters, the underlying sequence patterns differ. This structural distinction may be a key differentiator between the functions of these two classes of immune proteins.

## 2 Results

The IMGT-defined CDR loops (31) were extracted from each chain of the 270 high-quality TCR structures in STCRDab (STCRDab set; 30). This is more than double the numbers used in previous studies of TCR canonical forms, Al-Lazikani *et al.* (seven structures) and Klausen *et al.* (116 structures) (2, 24). Redundant sequences were retained as the conformations may differ as shown in the forthcoming sections. Summary statistics of the sequence-redundant STCRDab set for each of the CDR types are listed in Table 1.

**Table 1:**
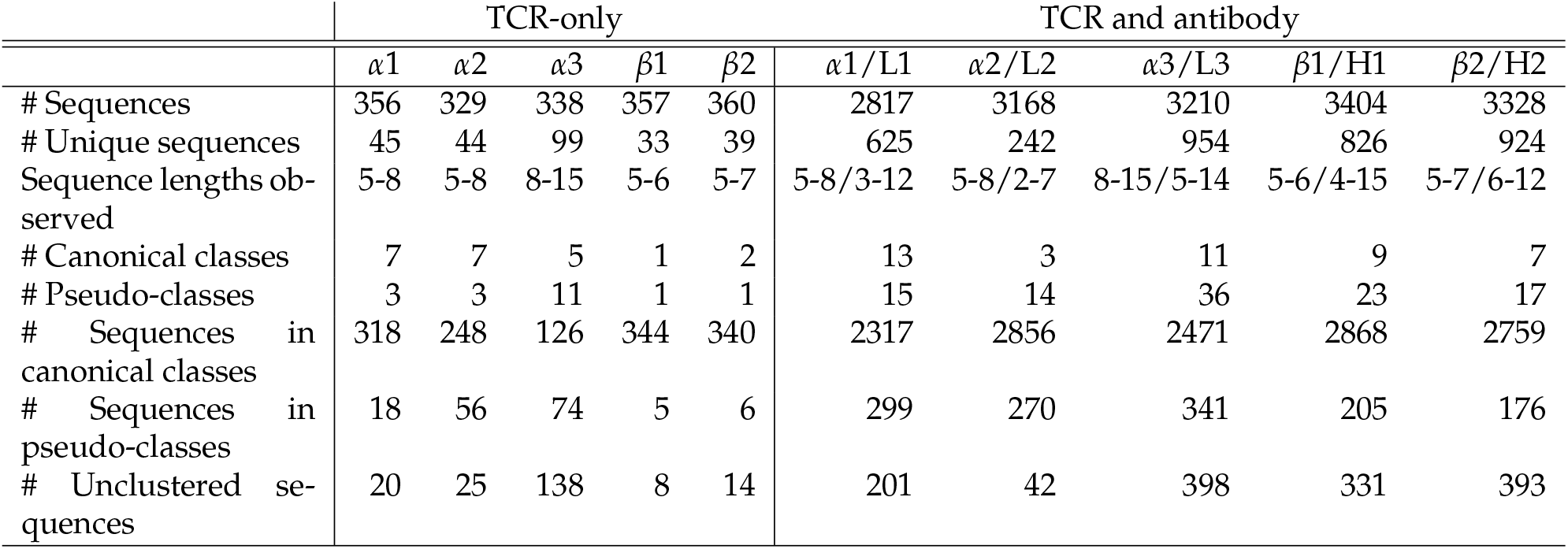
Summary of TCR and antibody CDR structural clusters. ‘#’ refers to the number of entities. TCR-only clusters are formed at a threshold of 1.0Å; TCR and antibody CDRs are clustered using the standard threshold for antibody CDRs given in Nowak et al. (40). The TCR and antibody CDR sequence lengths observed in the combined TCR and antibody CDR cluster are separated by ‘/’.

|  | TCR-only |  |  |  |  | TCR and antibody |  |  |  |  |
| --- | --- | --- | --- | --- | --- | --- | --- | --- | --- | --- |
| | $\alpha 1$ | $\alpha 2$ | $\alpha 3$ | $\beta 1$ | $\beta 2$ | $\alpha 1/L1$ | $\alpha 2/L2$ | $\alpha 3/L3$ | $\beta 1/H1$ | $\beta 2/H2$ |
| # Sequences | 356 | 329 | 338 | 357 | 360 | 2817 | 3168 | 3210 | 3404 | 3328 |
| # Unique sequences | 45 | 44 | 99 | 33 | 39 | 625 | 242 | 954 | 826 | 924 |
| Sequence lengths observed | 5-8 | 5-8 | 8-15 | 5-6 | 5-7 | 5-8/3-12 | 5-8/2-7 | 8-15/5-14 | 5-6/4-15 | 5-7/6-12 |
| # Canonical classes | 7 | 7 | 5 | 1 | 2 | 13 | 3 | 11 | 9 | 7 |
| # Pseudo-classes | 3 | 3 | 11 | 1 | 1 | 15 | 14 | 36 | 23 | 17 |
| # Sequences in canonical classes | 318 | 248 | 126 | 344 | 340 | 2317 | 2856 | 2471 | 2868 | 2759 |
| # Sequences in pseudo-classes | 18 | 56 | 74 | 5 | 6 | 299 | 270 | 341 | 205 | 176 |
| # Unclustered sequences | 20 | 25 | 138 | 8 | 14 | 201 | 42 | 398 | 331 | 393 |

### 2.1 Updating the canonical classes of TCR CDRs

We clustered each CDR type (*e.g.* CDR*α*1, CDR*α*2, etc.) using the DBSCAN method (40), with a minimum cluster size offive and a clustering threshold of 1.0 Å. For clusters with more than two unique sequences, we designated them as ‘canonical classes’; otherwise, they were considered ‘pseudo-classes’ (Table 1; see Materials and Methods for full details).

In total, we identified seven *α*1, seven *α*2, five *α*3, one *β*1 and two *β*2 canonical classes (see Table 1). The representative structures and sequence patterns of our TCR CDR canonical classes are shown in Figures 2 and S1 – S4. We found that some canonical forms have highly conserved positions in their encoding sequences, which might govern the conformations. However, we also observed many cases where structures of the same sequence fell into different structural clusters. For instance, the *α*1 loop with the sequence DSVNN belonged to *α*1-5-A when unbound, to pseudo-class *α*1-5-B* when bound to an MHC with a long peptide, and was unclustered when the binding peptide is short (Figure S5).

**Figure 2:**
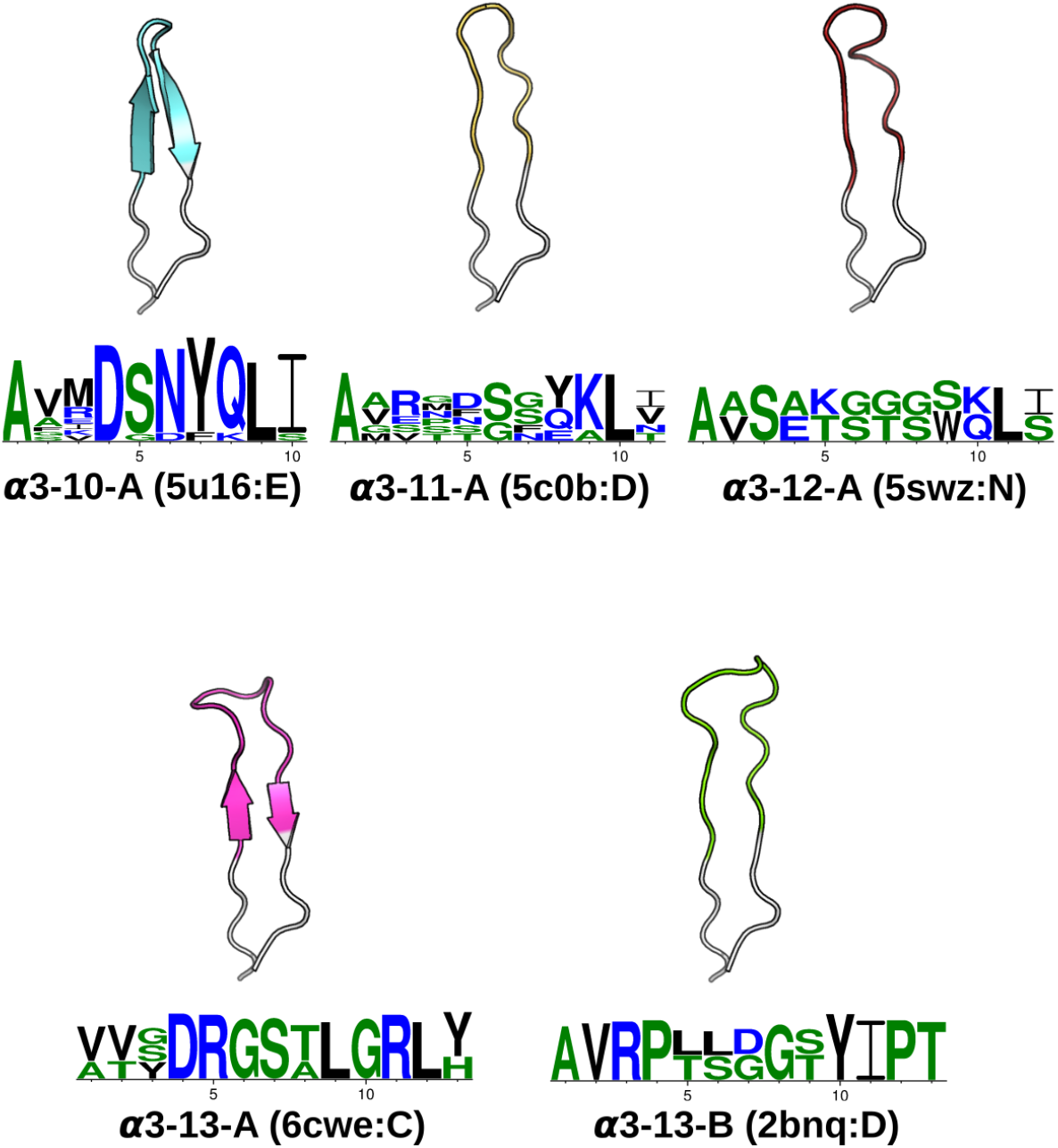
CDR*α*3 canonical classes. At a 1.0Å clustering threshold, our DBSCAN method identified five canonical classes. Each class has at least five structures and two unique sequences. For every CDR*α*3 class, the centroid structure is illustrated, with anchors in white, and the CDR*α*3 region (IMGT 105–117) coloured. The Protein Data Bank (PDB; 7) four-letter code and the chain identifier of the centroid structure is shown in the bracket next to the cluster name. The sequence pattern below each centroid structure is generated by WebLogo (12), using the unique sequences of the cluster: *α*3-10-A has 11,*α*3-11-A has 6,*α*3-12-A has 2,*α*3-13-A has 3 and *α*3-13-B has 2.

We compared our canonical forms to those from Al-Lazikani *et al.* and Klausen *et al.* (see Tables S1, S2). We matched canonical classes if their representative structure was found in our canonical class. Some canonical classes were represented by structures that were filtered out of our dataset for quality reasons. In these cases, a sequence-identical CDR with a comparable back-bone conformation was used as a proxy to match canonical classes (see Materials and Methods).

#### 2.1.1 Comparison to previous canonical forms

For CDR*α*1, our DBSCAN method broadly agreed with Al-Lazikani *et al.* and Klausen *et al.*(2, 24), apart from *α*1-3 in Al-Lazikani *et al.*’s classes, for which we found no corresponding cluster. Klausen *et al.*’s *α*1-5 cluster was matched to pseudo-class *α*1-6-F, meaning that for this cluster, we found more than five structures, but only a single unique sequence.

All seven of our CDR*α*2 classes and our one CDR*α*2 pseudo-class were matched to previously observed CDR*α*2 canonical classes (see Table S1 for the full list of comparisons). All five of our CDR*α*3 canonical forms and two of the pseudo-classes mapped to ones from Klausen *et al.*. In addition, we found nine further pseudo-classes that were not identified in their study. The cluster representative of the Klausen *et al.*’s *α*3-1 canonical class was filtered out of our dataset as it comes from a structure with a resolution greater than 2.9Å. Since its sequence was also absent in the rest of our dataset, we were unable to map this class to our canonical forms. The cluster representatives of Klausen *et al.*’s *α*3-2,*α*3-4,*α*3-11 and *α*3-12 were unclustered in our analysis.

Our single CDR*β*1 class and our one CDR*β*1 pseudo-class were matched to previous clusterings. Klausen *et al.*’s *β*1-3 and *β*1-4 forms were not in clusters in our work (Table S2). For CDR*β*2, we were unable to find a match for Al-Lazikani *et al.*’s *β*2-2 class, nor Klausen *et al.*’s *β*2-1 and *β*2-7 classes. However, our two CDR*β*2 classes and one CDR*β*2 pseudo-class were matched, with our *β*2-6-B merging three of Klausen *et al.*’s clusters (Table S2).

Both previous clusterings were based on backbone dihedral angles of the CDR loops, whereas in our work, we clustered using backbone distances. Despite these different approaches, there was a large degree of overlap between our canonical forms and those found previously. We have also identified a small number of new canonical classes from our larger dataset. As was shown for antibody CDR canonical forms (55), the growth of structural data continuously modifies our understanding of CDR loop structures. It is therefore necessary to continuously and preferably automatically update the definition of canonical forms as more structural information becomes available.

### 2.2 Prediction of CDRs from sequence

As our TCR canonical classes showed conserved sequence patterns (Figure 2 and S1 – S4), we built a sequence-based, length-independent position-specific scoring matrix (PSSM), that can be used to predict TCR CDR canonical classes (55). The performance of the predictor was evaluated by a leave-one-out cross-validation protocol on the unique sequences in STCRDab set (Table 2). Accuracy ranged from 73.2% to 100%, which is comparable to previous results (24).

**Table 2:** Leave-one-out cross-validation accuracy

| CDR | Unique sequences | PSSM Accuracy |
| --- | --- | --- |
| CDRα1 | 44 | 81.8% |
| CDRα2 | 43 | 73.2% |
| CDRα3 | 91 | 76.1% |
| CDRβ1 | 32 | 100% |
| CDRβ2 | 38 | 92.1% |

To assess the coverage of our method on large sets of sequencing data, we used our PSSMs to predict the CDR canonical classes of an NGS dataset of mouse TCR*α* sequences (32). The entire dataset contained 1,563,876 sequences (1,498,254 CDR*α*1, 1,563,876 CDR*α*2, 1,267,235 CDR*α*3); on a single 3.4GHz core, the prediction took three minutes. Our method achieved high coverage for CDR*α*1 and CDR*α*2, but made predictions for only 37% of the non-redundant CDR*α*3 sequences (Table 3). The poor coverage of CDR*α*3 could be due to the paucity of the currently available data in capturing the conformations of this more sequence-diverse CDR.

**Table 3:** Prediction of CDR*α* sequences from Tfh and Tfr cells.

| CDR | Prediction of redundant Sequences | Prediction of unique sequences |
| --- | --- | --- |
| CDRα1 | 283940/292310 | 1084/1278 |
| CDRα2 | 313317/313546 | 1359/1435 |
| CDRα3 | 116246/232260 | 1520/4139 |

Ten TCR structures containing 44 individual chains that were unseen at the time of methodology development, were used as a blind test set for the predictor. All canonical CDRs had 100% prediction coverage. Apart from CDR*α*3, all CDR types were predicted with 100% accuracy; in other words, at least one member of the predicted canonical class had backbone root-mean square deviation (RMSD) ≤1.0Å to the native structure. CDR*α*3 had one false prediction where the loop GTERSGGYQKVT was not assigned to any clusters even though the backbone RMSD falls within 1.0Å to a member of the *α*3-9-A canonical class.

### 2.3 Comparison between TCR and antibody CDRs

Inspired by the shared architecture and genetic generation mechanism of antibodies and TCRs, we examined whether their CDR length distributions, structures/canonical forms, and sequence patterns that give rise to their canonical forms overlap. For each of the TCR CDRs, we compared all its structures with the corresponding antibody CDR’s structures (*e.g.* CDR*α*1 with CDRL1). We considered the CDR sequence length distributions and their structural clustering with DBSCAN, using the clustering thresholds that have been previously used for antibody CDR clustering (40). Changing these thresholds does not qualitatively affect the results described below. A full list of comparisons is described in Table 1. For all pairs of TCR/antibody CDRs, we found that most clusters contained only TCR CDRs or antibody CDRs (*e.g.* CDR*α*1 only). In other words, the CDRs from TCRs and from antibodies tended to occupy distinct areas of structural space. Where we observed an overlap in the structural space, we inspected whether the sequence motifs for the canonical classes differ between the two types of receptors.

#### 2.3.1 CDR*α*1/CDRL1

The CDR*α*1 loops in our set were five-to seven-residues long, while CDRL1 loops spanned from three-to twelve-residues (Figure S9). The most common length was six for both types of loops: over 70% of the CDR*α*1 loops and 45% of the CDRL1 loops. Thirteen structural clusters of CDR*α*1/CDRL1 were identified (Figure S10), of these only *α*1,L1-5-A contained both CDR*α*1 loops (6 unique sequences) and CDRL1s (18 unique sequences). The sequence logos formed by the sequence-unique CDR*α*1 and CDRL1 loops in *α*1,L1-5-A had different sequence patterns (Figure S11). The general physicochemical properties were similar, but CDRL1 had a preference for Valine and Tyrosine on the third and fifth positions while CDR*α*1 showed ambiguity at these two positions. Principal component analysis (PCA) on one-hot-encoded unique sequences displayed a separation between CDR loops from the two types of receptors (see Figure S11), except for one TCR *α*1 sequence (DSVNN) that was considered to have a more similar sequence pattern to antibody L1 loops. This sequence is a TCR *α*1 sequence that adopts multiple conformations as described above.

#### 2.3.2 CDR*α*2/CDRL2

CDR*α*2 loops tend to be longer than CDRL2, with nearly all of the CDRL2 loops being three-residues long and CDR*α*2 having a range of lengths from five to eight (Figure S12). Given this, there is little chance of structural similarity between the two types of loops. Structural clustering confirmed that all classes only contained one CDR type, with two CDR*α*2 clusters, and one CDRL2 cluster (Figure S13).

#### 2.3.3 CDR*α*3/CDRL3

The majority of the CDRL3 loops in our set were nine-residues long, whilst CDR*α*3 loops had nine to fifteen residues (Figure 3). Among the eleven clusters identified, we found a cluster, *α*3,L3-10-A, which included both CDR types, with 17 and 12 unique sequences of CDRL3 and CDR*α*3 respectively (Figure 4). A PCA of the sequences in this cluster (*α*3,L3-10-A) separated the TCR*α*3 and antibody L3 members with one exception. The *α*3 sequence GTYNQGGKLI clustered with the L3 group. This sequence was one of the eight (out of a total of 99) unique *α*3 sequences we found to have multiple conformations. Our dataset contained three structures of this sequence; the other two were unclustered (3vxu:I and 3vxu:D).

**Figure 3:**
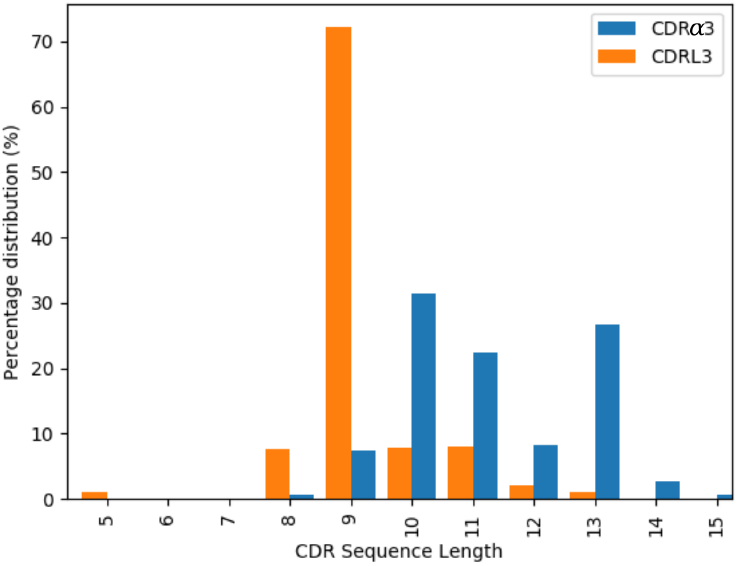
Length distributions of CDR*α*3 (blue) and CDRL3 (orange) loops.

**Figure 4:**
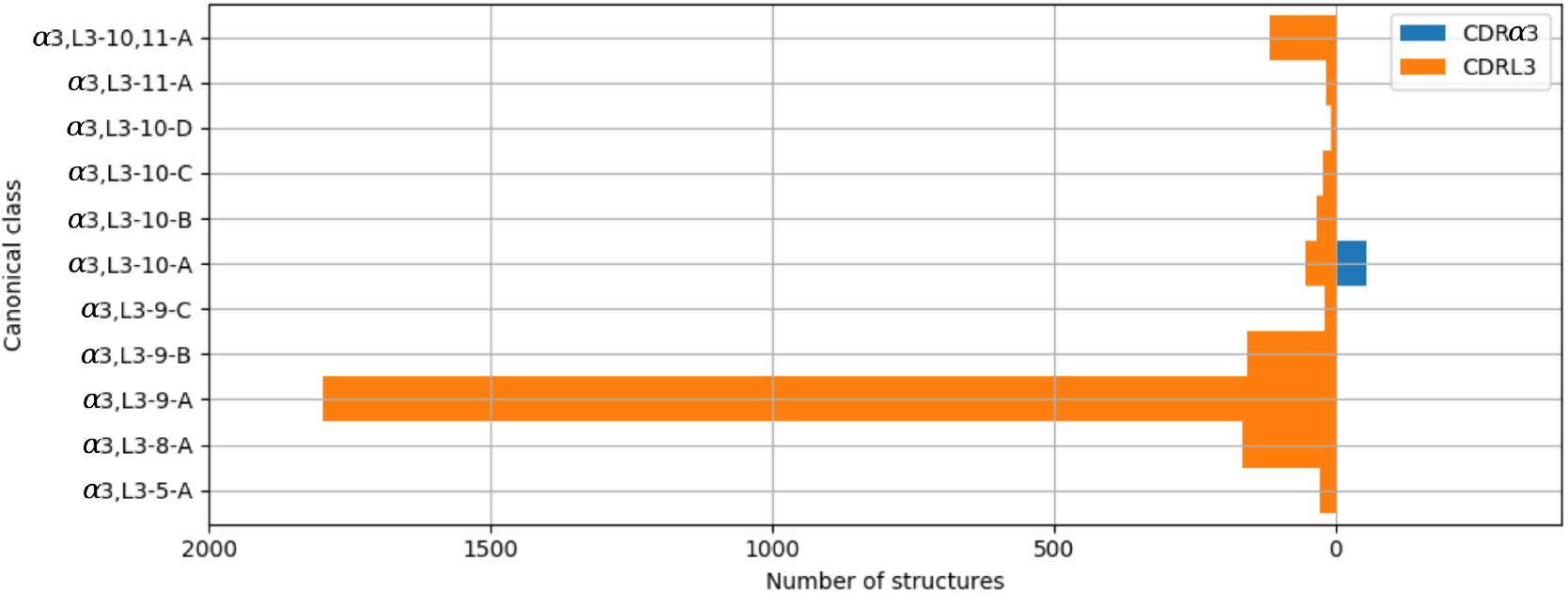
All CDR*α*3 and CDRL3 structures were clustered using the DBSCAN method. Orange bars indicate the number of CDRL3 structures, and blue bars the number of CDR*α*3 structures. All classes, apart from *α*3,L3-10-A, have structures from only one CDR type, *i*.*e*. CDR*α*3 or CDRL3.

#### 2.3.4 CDR*β*1/CDRH1

In our dataset, CDR*β*1 loops tended to be shorter than their CDRH1 counterpart (Figure S14). Over 80% of the CDR*β*1 were five-residues long, compared to eight-residues long loops dominating for CDRH1 (*>*80%). Nine classes were formed by the structural clustering: seven CDRH1 clusters and two CDR*β*1 clusters (Figure S15).

#### 2.3.5 CDR*β*2/CDRH2

More than 90% of the CDR*β*2 loops in our set were six-residues long, while only 0.2% of the CDRH2 had six residues and the rest were in the range of seven to ten residues (Figure S16). None of the seven clusters that were formed contained members from both types of CDRs (Figure S17).

#### 2.3.6 *β*-chain CDRs in TCRs and heavy-chain CDRs in antibodies and nanobodies

In our clustering, CDRs from nanobodies were also included. Unlike the CDRs of TCRs, these typically did fall into the antibody H1 and H2 clusters (Figures S15, S17). In the case of CDR*β*1/CDRH1, the majority of the nanobody CDRH1 loops were clustered with antibody CDRH1s in the *β*1,H1-8-A class. The remaining nanobody CDRH1 loops formed the majority of the other two length-eight clusters. Nanobody CDRH2 loops clustered with the length-seven and eight antibody CDRH2 loops. These results suggest that nanobody CDR loops are structurally more similar to antibodies than to TCRs.

**Figure 5:**
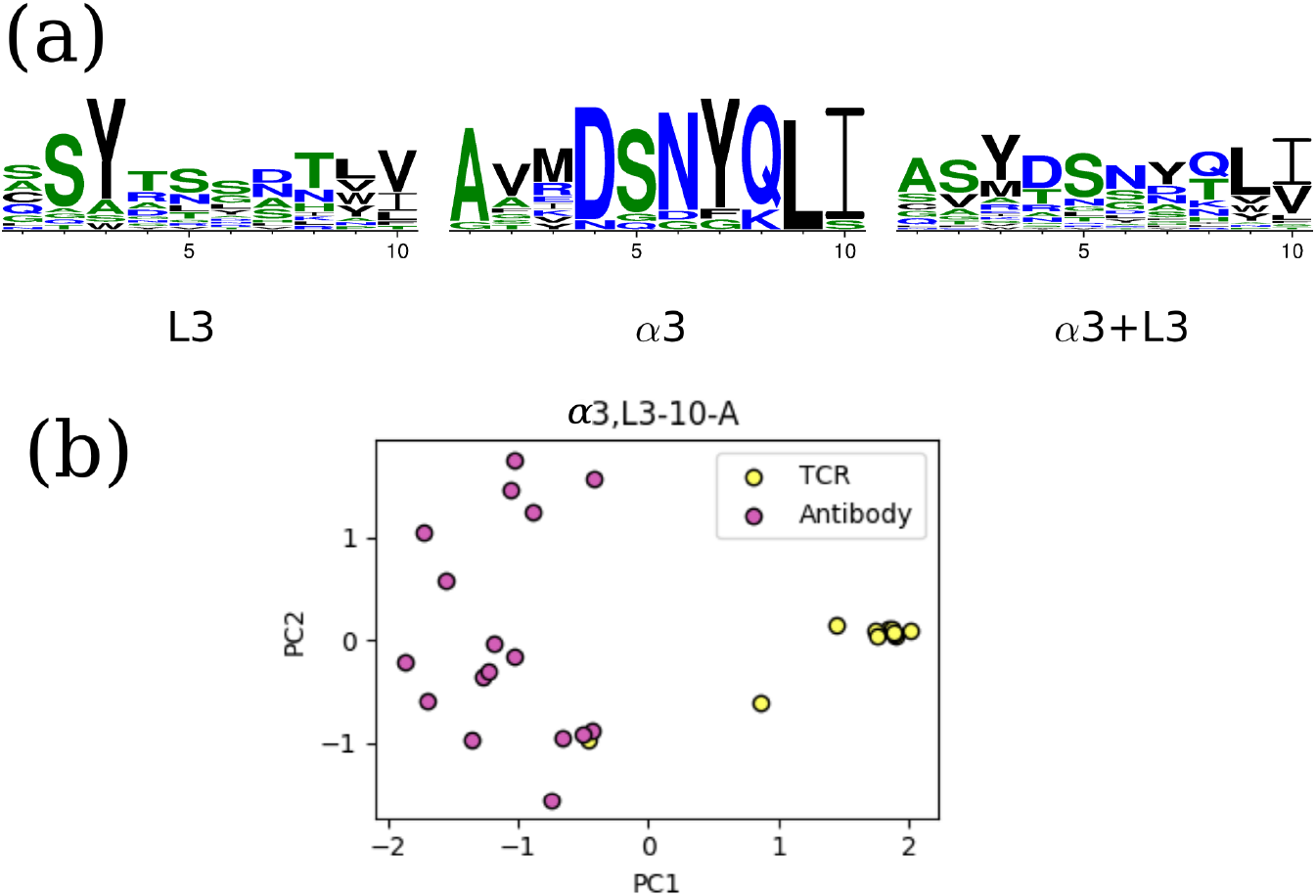
The unique TCR and antibody sequences in the *α*3,L3-10-A class. (a) Sequence logos of CDRL3 and CDR*α*3 loops in the *α*3,L3-10-A class, using only the unique sequences. The sequence patterns appear to be distinct between the two classes. (b) Principal component analysis (PCA) plot of the first two components in one-hot-encoded unique sequences in *α*3,L3-10-A, stratified by TCR and antibody CDRs (see Materials and Methods). TCR *α*3 and antibody L3 are separated by the first principal component, only one TCR sequence (GTYNQGGKLI) is close to the antibody set.

#### 2.3.7 CDR*β*3/CDRH3

There are no canonical classes for CDRH3 or CDR*β*3 but in the case of CDRH3, previous studies have found structural conservation in the start and end of the loop, known as the ‘torso’ or ‘base’ region (19, 54). We therefore inspected the base structure of the CDR*β*3/CDRH3 loops. Following the study by Weitzner et al. (54), we carried out the loop anchor transform (LAT) analysis that was used to capture the characteristic base structure in CDRH3. The LAT analysis showed that CDR*β*3 loops have a similar width of distribution in all six degrees of freedom as seen for CDRH3, but with a slight shift in the peak (Figure S18). This presented the possibility that CDR*β*3 and CDRH3 could share similar base structures. In the same study, the pseudo bond angle (*τ*) and pseudo dihedral angle (*α*) of the penultimate residue of the CDRH3 structure (IMGT position 116) were used to differentiate between extended and kinked torsos. We found that very few CDR*β*3 loops had their *τ* _116_ and *α* _116_ in the space of kinked torsos (Figure 6). Instead the majority of CDR*β*3 loops had an extended base. This behaviour is the opposite of CDRH3 loops. The eight outlying CDR*β*3 structures with positive *α* _116_ have either an aromatic side chain at IMGT position 116 that restricts the shape of the base, or an abrupt bend to accommodate unusual binding peptides. Consistent observations were made when we analysed the *ϕ*/*ψ* plots for the torso positions as outlined in Finn et al. (19, see Figure S19).

**Figure 6:**
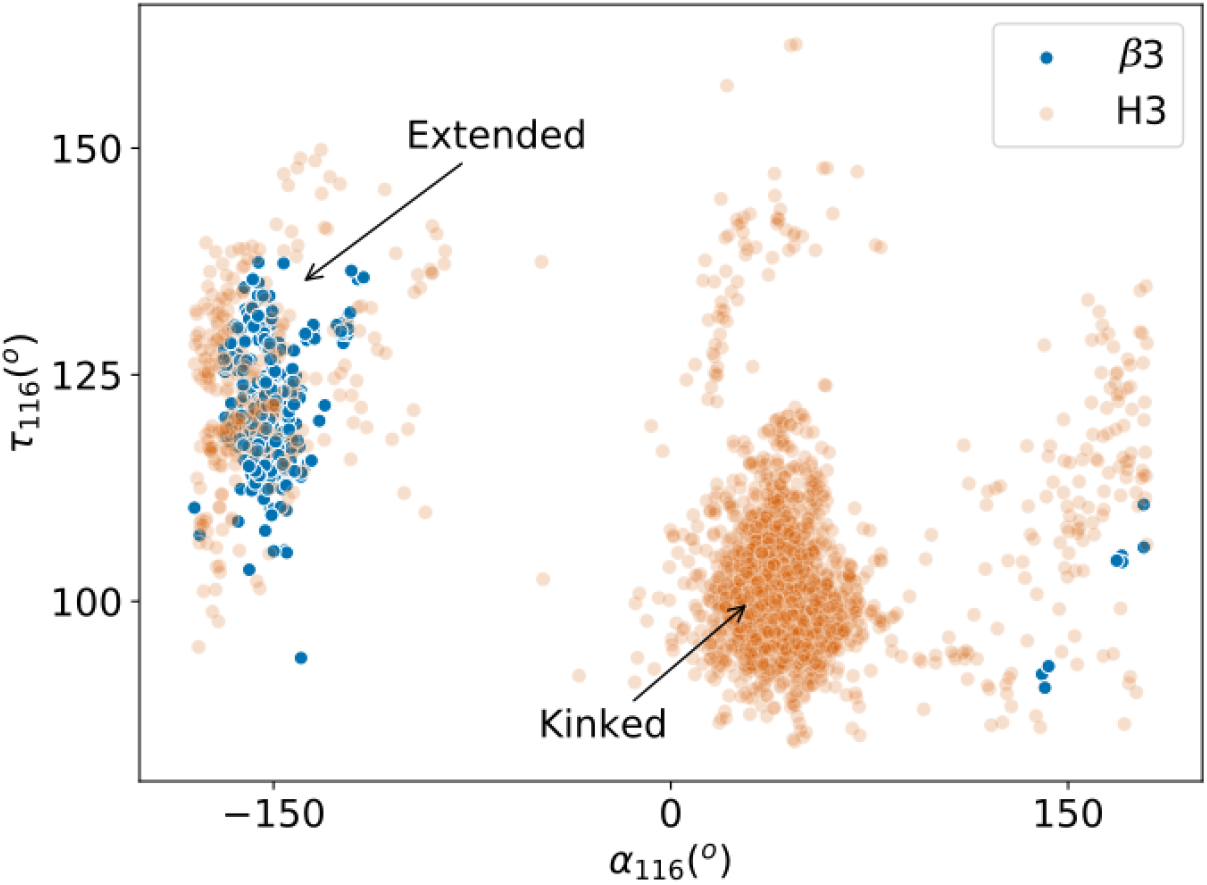
Pseudo bond angle (*τ*) and pseudo dihedral angle (*α*) analyses on IMGT (31) position 116. Scatters represent individual observations of *β*3 (blue) and H3 (orange) structures. Regions occupied by kinked and extended CDRH3 loops according to Weitzner et al. (54) are indicated by arrows.

#### 2.3.8 CDR structural variability in TCR and antibodies

As noted above, some TCR CDRs with identical sequences fell into different structural clusters. Considering the joint clustering of TCR and antibody CDRs, we found that of the 212 unique TCR CDR sequences with multiple example structures, 41 (19.3%) fell into more than one structural cluster (Table S3). This happened far less for antibody CDRs where only 7.95% (166 of 2089) of the unique antibody CDR sequences with multiple example structures were found in more than one structural cluster. Whether the structure was crystallised in complex with the cognate molecule did not appear to cause the conformational difference as bound and unbound structures were observed in the different clusters (Tables S4, S5).

Following the analysis in (35), we compared the maximum backbone RMSD of structures from an identical sequence (Figure S20). If we consider structures with a difference of less than 1Å in backbone RMSD as similar (184 of the 284 TCRs and 2195 of the 2717 antibodies), this analysis illustrates that the majority of the sequence-identical structures are structurally similar in both TCR and antibody CDRs, but that TCR loops show a more skewed distribution, tending towards higher structural variability.

## 3 Discussion

We have identified and refined the canonical class definitions of TCR CDRs using a more up-to-date structural dataset, and used these to generate an auto-updating database and prediction server.

In our dataset of 270 TCR structures, we find seven CDR*α*1, seven CDR*α*2,five CDR*α*3, one CDR*β*1, and two CDR*β*2 canonical classes. In addition, we report several ‘pseudo-classes’, in which there are multiple examples of the structural conformation, but only one unique sequence. One of the major advantages of a canonical class model is the rapid mapping between the sequence and structural space. To demonstrate this for TCRs, we applied an adapted PSSM-based methodology (55) to assign the canonical classes for an NGS dataset of ~1.5 million mouse TCR*α* sequences in a few minutes.

The commonalities between TCR and antibody in their folds and the genetic mechanisms that generate them prompted us to explore the similarities and differences between TCR and antibody CDR structures. We performed a length-independent structural clustering of TCR and antibody CDRs and found that they almost always separate into different clusters. This separation was partly driven by the differences in the length distributions between TCR and antibody CDRs. In the cases where we found structural clusters which contained both types of loops, we found that the underlying sequence motifs were distinct.

We also compared the CDR*β*3 and CDRH3 loops in terms of their torso structures – the starting and the ending residues of the CDR*β*3/CDRH3 loops, and found once again TCRs and antibodies gravitated towards different structures. CDRH3 loops in antibodies tend to have a kinked torso while CDR*β*3 are only found with the extended torsos that is less common in CDRH3.

Driven by the multiple observations where sequence-identical TCR CDR structures were found in different clusters, we also assessed the structural variability of TCR and antibody CDRs. Our results agree with previous findings about the flexibility of TCR CDRs (13, 22), and further suggest that TCR loops are more flexible than antibody CDRs. Structural rigidification has been proposed as an affinity maturation mechanism in antibodies (37), while flexibility in the binding site has been suggested as one of the features enabling the promiscuous binding of TCR-MHC (43). The fact that nearly 20% of the TCR CDRs that could show different conformations did so suggests that the canonical class model will struggle to accurately predict TCR CDR conformations. Overall, our results suggest that there are structural differences between TCR and antibody CDRs, and the differences we observe potentially help explain how these two receptors bind to their separate target antigens.

Many groups have attempted to augment TCR and antibody designs by swapping binding sites between these two receptors (*e.g.* 56, 57). TCR-mimic antibodies were developed in the hope of transferring the ability of TCRs to target intracellular proteins, to antibodies for the use in cancer therapy. Antibody-like TCR design transfers the highly specific antibody binding sites to TCRs. It was shown to enhance the specificity and affinity in TCR antigen recognition *in vitro* and *in vivo*(57). These cases present the possibility of altering the behaviour of the receptors such as targeting peptides in the context of an MHC molecule by grafting TCR binding sites, or enhancing specificity when the antibody components are added. Our results should aid the development and design of these types of therapeutic immune proteins.

## 4 Materials and Methods

### 4.1 Nomenclature

We use the following nomenclature for structures: four characters of the PDB code, followed by a colon, then the chain identifier of the structure, *e.g.* 5hhm:E. Clusters are identified by two letters describing the CDR type, the loop length(s), then a letter (40). For instance, the *α*1-6-B class refers to the length-6 CDR*α*1 class with the second-largest number of unique sequences. Pseudo-clusters with only one unique sequence have their names appended with an asterisk (*). For the clustering of both TCR and antibody CDRs, we name the clusters by the two letters representing the TCR CDR type, two letters for the antibody CDR type, the loop length(s), followed by a letter, *e.g.α*1,L1-11,12-A denotes the largest cluster from the CDR*α*1/CDRL1 clustering, in which sequences are 11 or 12 residues long.

### 4.2 Definitions

TCR and antibody structures were numbered in the IMGT scheme (31) using ANARCI (15): CDR1 (27–38), CDR2 (56–65), and CDR3 (105–117). To compare the CDR loops of TCRs and antibodies, we assumed equivalence between *β*/heavy chains and *α*/light chains (30), as TCR*β* and antibody heavy chains rely on *VDJ* recombination, while TCR*α* and antibody light chains are formed by *VJ* recombination.

### 4.3 Structural datasets

TCR structures with at least one *α* or *β* chain and resolution ≤2.8Å were downloaded from STCRDab on 31 May 2018 (30). Antibody structures with resolution ≤2.8Å were downloaded from SAbDab on 31 May 2018 (17). In total, 270 TCR PDB entries and 2563 antibody PDB entries were used. We retained all chains in a PDB entry that passed the quality criteria. The structures of the CDR loops were extracted using a similar procedure to Nowak *et al.*(40). The IMGT-defined CDRs, along withfive N-terminal andfive C-terminal anchor residues, were selected if there were no missing backbone atoms and none of the backbone atoms had B-factors *>*80. Loops also needed to be continuous, *i.e.*, all peptide bond lengths were *<*1.37Å. Since loops with identical sequences can adopt multiple conformations (see Results and Figure S5 in the current work on TCR CDRs and examples in (40) for antibody CDRs), we retained all CDR loop structures that passed the structural quality criteria.

### 4.4 Clustering method

Loops are clustered using the length-independent clustering method from Nowak *et al.*(40). First, the algorithm superimposes the backbone atoms of all ten anchor residues. Structural similarity is then calculated using the dynamic time warp (DTW) algorithm (47); the resulting DTW score is effectively a length-independent RMSD value (40). The DTW scores are then clustered using the DBSCAN method (18).

In the clustering of TCR CDRs alone, we only consider a set of sequences to be a CDR cluster only if it contains a minimum of five structures and two unique sequences. If a set contains five or more structures but only one unique sequence, we label this a ‘pseudo-class’. All other loops are considered to be ‘unclustered’. This is a more lenient threshold than previous investigations clustering antibody CDR loops (*e.g.*40), as the dataset of TCR structures is far smaller. To choose the optimal clustering parameter, DBSCAN was run over a range of DTW score thresholds in increments of 0.1Å. We find that 1.0 Å offered an optimal balance between the number and size of clusters for the five TCR CDR loops.

For the clustering of multiple CDR types, we use the same clustering thresholds and criteria for the respective CDR types as the antibody study (40), where they were selected using the Ordering Points to Identify the Clustering Structure (OPTICS; 4) algorithm. CDR*α*1/CDRL1 are clustered at 0.82Å, CDR*α*2/CDRL2 at 1Å (not initialised in the previous paper), CDR*α*3/CDRL3 at 0.91Å, CDR*β*1/CDRH1 at 0.8Å and CDR*β*2/CDRH2 at 0.63Å. A valid cluster must contain at least six unique sequences as illustrated in Nowak et al. (40).

### 4.5 Comparison with TCR canonical classes in earlier work

In order to compare our CDR clusters to previous studies (2, 24), we identified overlaps in the representative PDB entries. For example, Al-Lazikani *et al.*’s *α*2-3 class contains the PDB entry 1tcr. Since 1tcr:A was in our *α*2-8-B class, we considered these two classes (*α*2-3 and *α*2-8-B) to be analogous. A similar procedure was used to map our classes to Klausen *et al.*’s classes (24).

Some canonical classes from Al-Lazikani *et al.* or Klausen *et al.* were represented by structures that were filtered out of our dataset. To match these classes, we searched for a CDR structure in our set that has the same CDR sequence and checked if there was a backbone match. For example, the centroid structure of Klausen *et al.*’s *α*3-7 class is PDB 3tf7, which has the sequence AVSAKGTGSKLS. We found that 3tfk:C has the same CDR*α*3 sequence as 3tf7 and a backbone RMSD of 0.81Å; thus, we assigned their *α*3-7 to our *α*3-12-A class.

### 4.6 Sequence-based prediction of canonical forms

For each CDR canonical class, we generated a position-specific scoring matrix (PSSM). The score of an amino acid *a* in IMGT position *i*,*s*(*a*,*i*), is

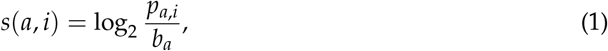

where *p*_*a,i*_ represents the probability of *a* at *i*, and this is calculated separately for each class. The background probability of *a*, *b*_*a*_, was assumed to be identical for all residues *i.e.* 0.05.

To predict the cluster for a target loop with length *l*, we first select PSSMs containing loops with the same length. For example, if the new CDR*α*1 loop is six residues long, we choose PSSMs of the *α*1-6-A, *α*1-6-B, *α*1-6-C and *α*1-6-D classes. The PSSM score for the target loop for class *c*, *P*_*c*_, is the sum of the position-specific scores

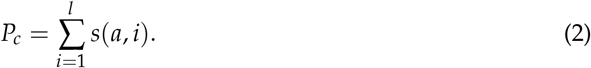

If *a* is never observed at *i*, we assume *s* (*a*,*i*) = −1. For canonical class assignment, we deignated a target loop to the class with the highest value of *P*_*c*_. Furthermore *P*_*c*_ must be higher than 1, except for CDR*α*3 where *P*_*c*_ must be equal to or greater than the loop’s length. The value of *P*_*c*_ was chosen by performing leave-one-out cross-validation tests over several values of *P*_*c*_. Sequences were assigned to a pseudo-class if and only if they had an identical sequence.

To benchmark the scoring strategy, we ran a leave-one-out cross-validation protocol on the unique sequences from canonical classes and unclustered structures. A prediction was evaluated using the following criteria:

- True positive: sequence is assigned to the correct canonical class.
- False positive: sequence is assigned to a different canonical class.
- True negative: sequence is from an unclustered loop, and not assigned a canonical class.
- False negative: sequence is from a canonical class, but predicted to be unclustered.

### 4.7 Prediction on next-generation sequencing dataset

We predicted the canonical forms for the *α*-chain in a set of mouse TCR sequences from BioProject accession PRJNA362309 (32). Overlapping Illumina reads were assembled using FLASh (33), and TCR amino acid sequences were extracted using IgBLAST (58). The sequences were then numbered by ANARCI (15); only those with productive CDR3 rearrangement, CDR1 and CDR2 loops at least five residues long, and CDR3 loops at least eight residues long were retained.

### 4.8 Prediction of new TCR structures

We used the 44*αβ* TCR structures that were released between 31st May 2018 and 5th June 2019 as a blind test set. Unlike our structural dataset for clustering, we did not impose a quality restriction for CDR prediction. Predictions were considered to be correct if the backbone RMSD between the native CDR structure and any member of the assigned canonical class was ≤1.0Å.

### 4.9 Comparison between TCR and antibody CDR structures

We clustered TCR and antibody CDR structures and examined the length distribution, structural clustering and sequence patterns. Sequence lengths of CDR loops disregard the five anchor residues at each side of the CDR structures. Structural clustering comparison were done as described above. We used WebLogo to generate all sequence patterns (12).

In canonical forms where both TCR and antibody CDRs were found, we examined the difference between the sequence patterns used by TCRs and antibodies. We transformed the sequences using one hot encoding by position, where each feature was represented by the position and the residue name, and applied Principal Component Analysis (PCA).

### 4.10 Analysis of CDR*β*3 and CDRH3 structures

We analysed CDR*β*3 structures with a set of metrics that have been previously applied to the CDRH3 loop in antibodies: loop anchor transform (LAT), pseudo bond angles and dihedral angles (19, 54).

LAT is the Euler transformation of the coordinate planes formed by residues at IMGT positions of 105 and 117 (see Supplementary Information of Weitzner et al. (54) for a detailed mathematical definition). Brie fly, a coordinate system is defined for each of the two residues centred on the C_*α*_ atoms, where the z-axis points towards the carbonyl carbon, the y-axis is perpendicular to z in the N-C_*α*_-C plane, and the x-axis is the cross product of these two components. The Euler transformation is then represented by six degrees of freedom, capturing the translation (*X*,*Y*,*Z*) and rotation (*ϕ*,*ψ*,*θ*).

We calculated the pseudo bond angle (*τ*) and pseudo dihedral angle (*α*) of the residue at IMGT position 116 (corresponding to Chothia position 101), in the CDR*β*3 and CDRH3 sets. Consistent with Weitzner et al. (54), the pseudo bond angle is formed by the C_*α*_ atoms of the residues before, at and after the IMGT position 116, whereas the pseudo dihedral angle spans across the C_*α*_ atoms of residues at position 116, one before and two after.

Finn et al. (19) observed that the dihedral angles adopted by the base residues of extended torsos were different from that of kinked torsos. To capture the observation made by Finn et al. (19), the dihedral angles of the first three (T1-T3) and last four (T4-T7) residues of the CDR*β*3 and CDRH3 loops are obtained from the Biopython module (10).

## Acknowledgement

We thank Aleksandr Kovaltsuk for his assistance in curating the sequencing datasets. This work was supported by the Engineering and Physical Sciences Research Council and Medical Research Council Grant EP/L016044/1.

## Notes

#### Summary of Updates

Removed first page of the Supplementary Materials as it overlaps with the last page of the main text.

