## Supplementary Materials for "Comparative analysis of the CDR loops of antigen receptors"

### 6 Supplementary Information

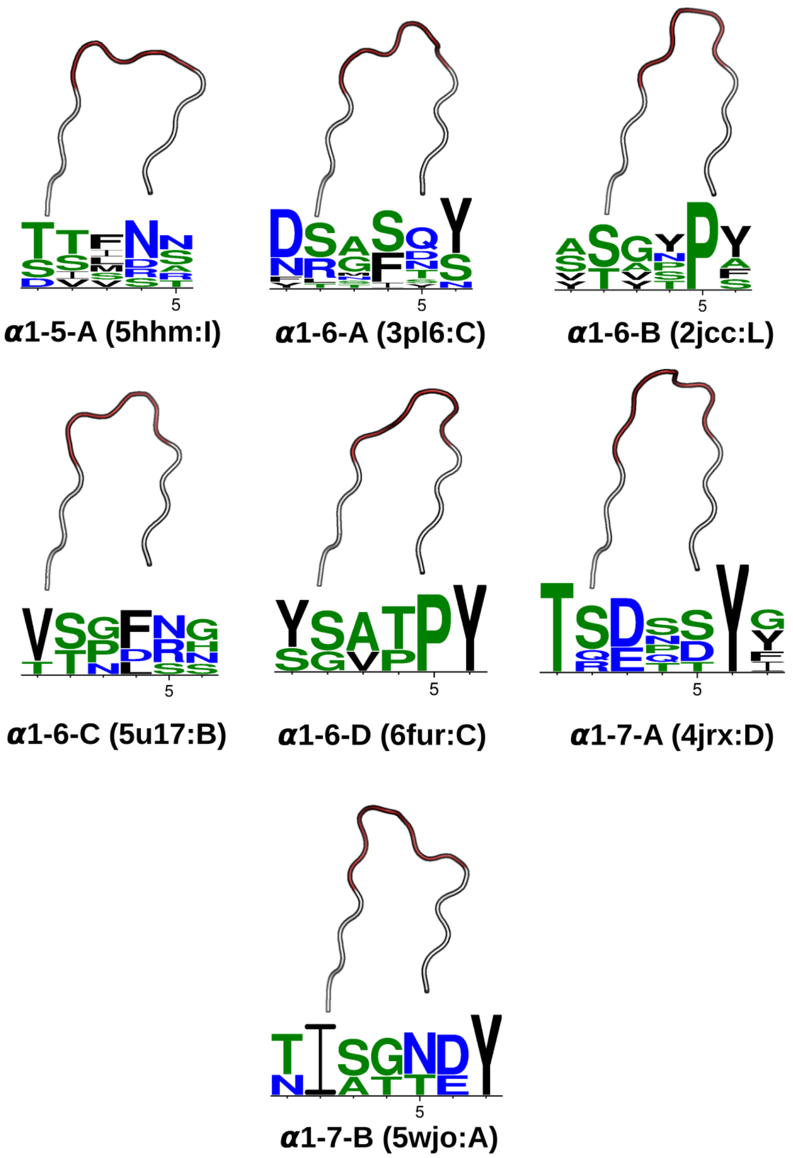


Figure S1: CDR$\alpha$1 loop clusters. At a 1.0Å clustering threshold, our DBSCAN method identified seven canonical classes (see Results). Each class has at least five structures and two unique sequences. Anchors are coloured white, while the CDR$\alpha$1 region (IMGT 27–38) is coloured. The Protein Data Bank (PDB; 7) four-letter code and the chain identifier of the centroid structure is shown in the bracket next to the cluster name. The sequence pattern below each centroid structure is generated by WebLogo (12), using the unique sequences of the cluster: $\alpha$1-5-A has 7, $\alpha$1-6-A has 13, $\alpha$1-6-B has 6, $\alpha$1-6-C has 5, $\alpha$1-6-D has 3, $\alpha$1-7-A has 6 and $\alpha$1-7-B has 3.

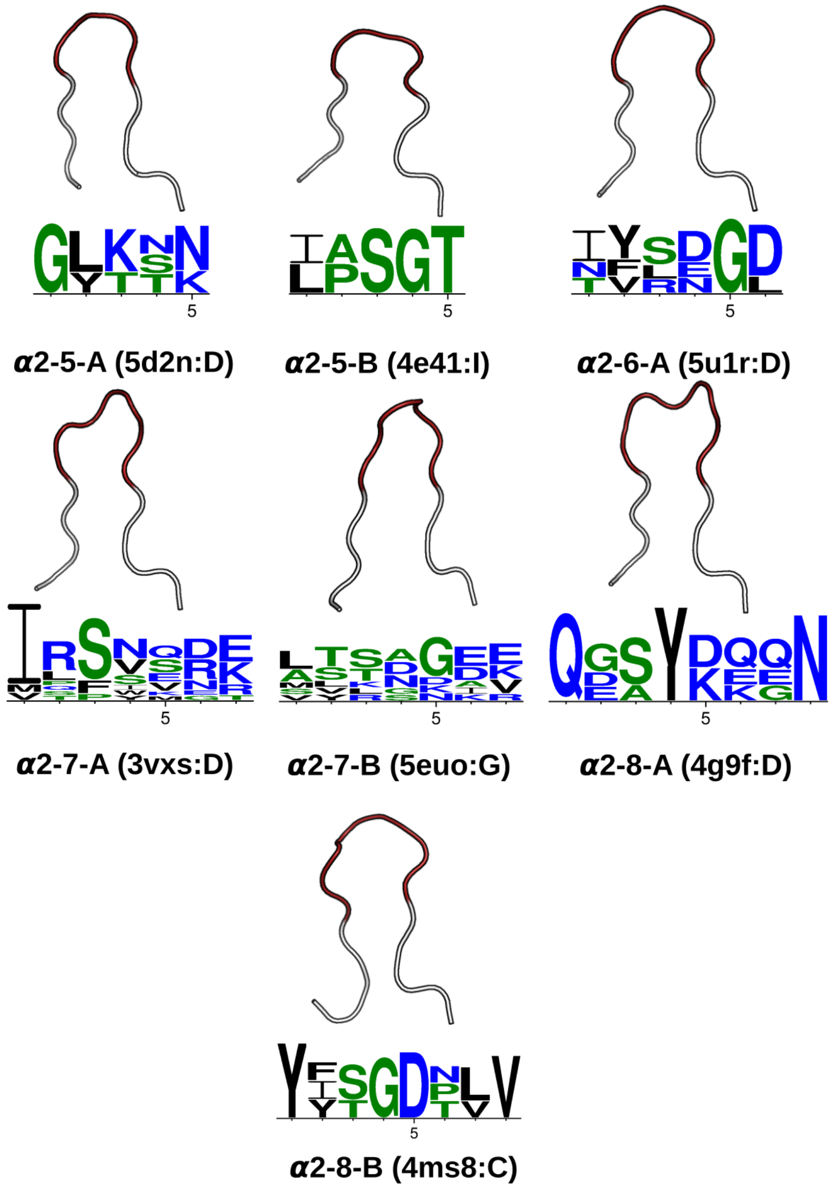


Figure S2: CDR$\alpha$2 loop clusters. At a 1.0Å clustering threshold, our DBSCAN method identified seven canonical classes (see Results). Each class has at least five structures and two unique sequences. Anchors are coloured white, while the CDR$\alpha$2 region (IMGT 56–65) is coloured. The PDB four-letter code and the chain identifier of the centroid structure is shown in the bracket next to the cluster name. The sequence pattern below each centroid structure is generated by WebLogo (12), using the unique sequences of the cluster: $\alpha$2-5-A has 3, $\alpha$2-5-B has 2, $\alpha$2-6-A has 4, $\alpha$2-7-A has 12, $\alpha$2-7-B has 8, $\alpha$2-8-A has 4 and $\alpha$2-8-B has 3.

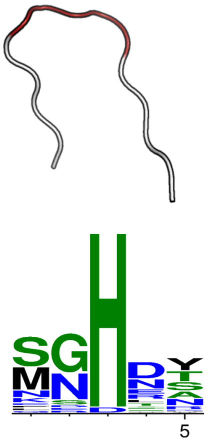

**β1-5-A (1bd2:E)**

Figure S3: CDRβ1 loop clusters. At a 1.0Å clustering threshold, our DBSCAN method identified one canonical class (see Results). Each class has at least five structures and two unique sequences. Anchors are coloured white, while the CDRβ1 region (IMGT 27–38) is coloured. The PDB four-letter code and the chain identifier of the centroid structure is shown in the bracket next to the cluster name. The sequence pattern below each centroid structure is generated by WebLogo (12), using the unique sequences of the cluster: β1-5-A has 30.

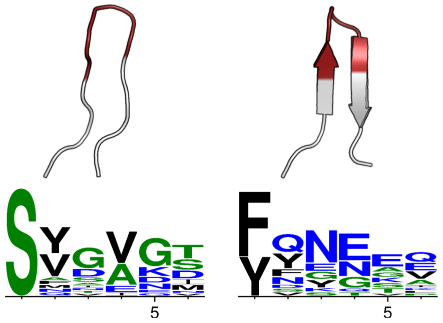

**β2-6-A (3qh3:B)** **β2-6-B (3vxm:E)**

Figure S4: CDRβ2 loop clusters. At a 1.0Å clustering threshold, our DBSCAN method identified two canonical classes (see Results). Each class has at least five structures and two unique sequences. Anchors are coloured white, while the CDRβ2 region (IMGT 56–65) is coloured. The PDB four-letter code and the chain identifier of the centroid structure is shown in the bracket next to the cluster name. The sequence pattern below each centroid structure is generated by WebLogo (12), using the unique sequences of the cluster: β2-6-A has 18 and β2-6-B has 16.

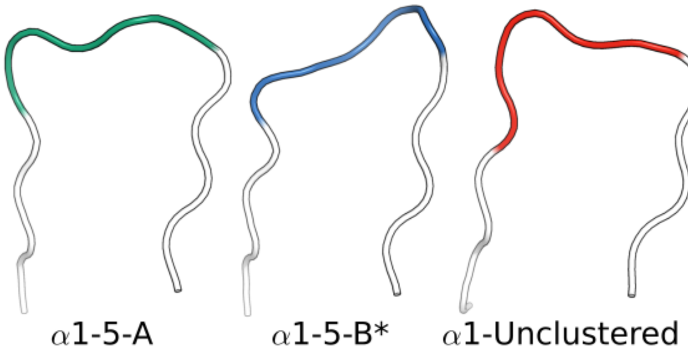


Figure S5: CDR$\alpha$1 loops with the sequence DSVNN can form the $\alpha$1-5-A canonical class (PDB: 5fka:A; unbound) or the $\alpha$1-5-B* pseudo-class (PDB: 2ian:D; bound to MHC with a long peptide). Alternatively, they can also be unclustered (PDB: 4mnq:D; bound to MHC with a short peptide).

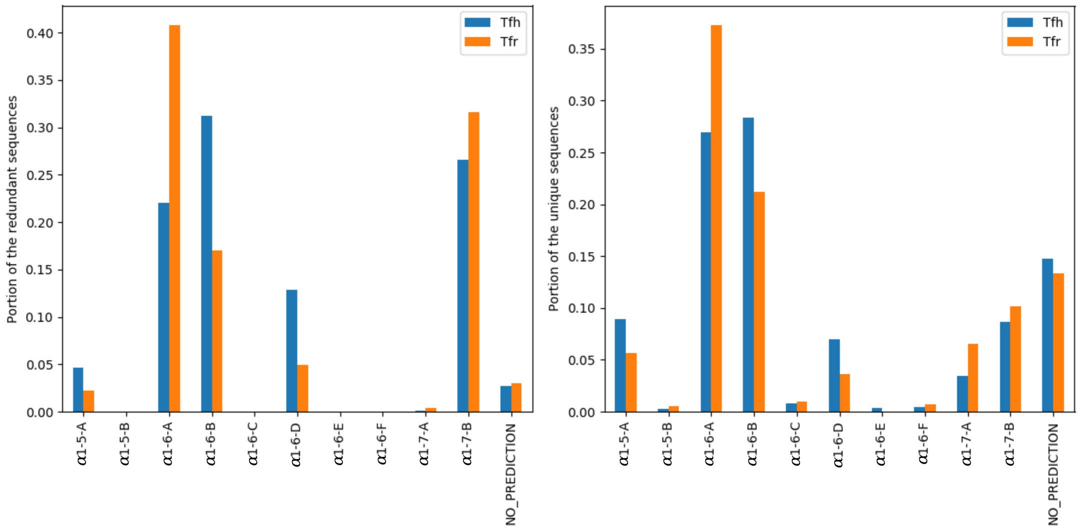


Figure S6: CDR$\alpha$1 canonical form prediction from the TCR sequences in Tfh and Tfr cells from (32).

| Canonical class | Al-Lazikani | Klausen | Number of unique sequences (Number of structures) |
|---|---|---|---|
| $\alpha$1-5-A | | $\alpha$1-1 | 7 (36) |
| $\alpha$1-5-B* | | | 1 (7) |
| $\alpha$1-6-A | $\alpha$1-1 | $\alpha$1-2 | 13 (106) |
| $\alpha$1-6-B | $\alpha$1-2 | $\alpha$1-3 | 6 (25) |
| $\alpha$1-6-C | | $\alpha$1-4 | 5 (93) |
| $\alpha$1-6-D | | | 3 (18) |
| $\alpha$1-6-E* | | | 1 (5) |
| $\alpha$1-6-F* | | $\alpha$1-5 | 1 (6) |
| $\alpha$1-7-A | | $\alpha$1-7 | 6 (16) |
| $\alpha$1-7-B | | $\alpha$1-6 | 3 (24) |
| (unclustered) | $\alpha$1-3 | | |
| $\alpha$1-unclustered | | | 10 (20) |
| $\alpha$2-5-A | | $\alpha$2-2 | 3 (16) |
| $\alpha$2-5-B | | $\alpha$2-1 | 2 (22) |
| $\alpha$2-6-A | $\alpha$2-1 | $\alpha$2-3 | 4 (94) |
| $\alpha$2-6-B* | | | 1 (9) |
| $\alpha$2-7-A | $\alpha$2-2 | $\alpha$2-4 | 12 (56) |
| $\alpha$2-7-B | $\alpha$2-4 | $\alpha$2-5 | 8 (41) |
| $\alpha$2-7-C* | | $\alpha$2-7 | 1 (28) |
| $\alpha$2-8-A | | $\alpha$2-8 | 4 (9) |
| $\alpha$2-8-B | $\alpha$2-3 | $\alpha$2-6 | 3 (10) |
| $\alpha$2-8-C* | | | 1 (19) |
| $\alpha$2-unclustered | | | 11 (25) |
| $\alpha$3-9-A* | | | 1 (5) |
| $\alpha$3-10-A | | $\alpha$3-3 | 11 (53) |
| $\alpha$3-10-B* | | | 1 (12) |
| $\alpha$3-10-C* | | | 1 (8) |
| $\alpha$3-10-D* | | | 1 (5) |
| $\alpha$3-10-E* | | | 1 (6) |
| $\alpha$3-11-A | | $\alpha$3-5 | 6 (27) |
| $\alpha$3-11-B* | | | 1 (7) |
| $\alpha$3-11-C* | | $\alpha$3-6 | 1 (9) |
| $\alpha$3-12-A | | $\alpha$3-7 | 2 (5) |
| $\alpha$3-12-B* | | | 1 (5) |
| $\alpha$3-13-A | | $\alpha$3-8 | 3 (33) |
| $\alpha$3-13-B | | $\alpha$3-10 | 2 (8) |
| $\alpha$3-13-C* | | $\alpha$3-9 | 1 (6) |
| $\alpha$3-13-D* | | | 1 (6) |
| $\alpha$3-13-E* | | | 1 (5) |
| (filtered) | | $\alpha$3-1 | |
| (unclustered) | | $\alpha$3-2/$\alpha$3-4/$\alpha$3-11/$\alpha$3-12 | |
| $\alpha$3-unclustered | | | 72 (138) |

Table S1: Summary of CDR$\alpha$ canonical classes. The CDR$\alpha$ canonical classes from our study are compared to the canonical classes of Al-Lazikani *et al.* (2) and Klausen *et al.* (24) (see Materials and Methods). Briefly, if there is at least one structure that is present in both our canonical class and those from the literature, we consider these classes to be analogous. Pseudo-classes are indicated with *.

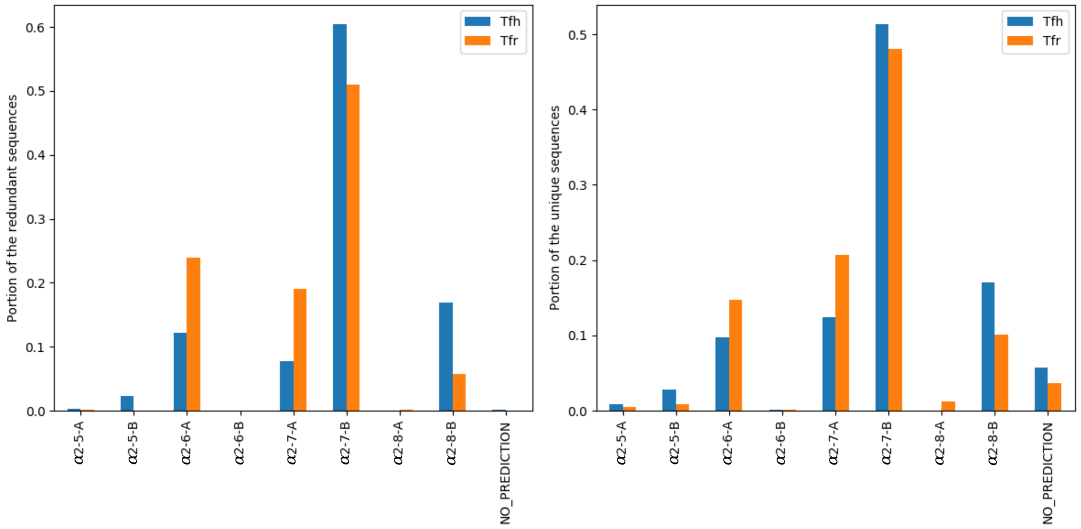


Figure S7: CDR$\alpha$2 canonical form prediction from the TCR sequences in Tfh and Tfr cells from (32).

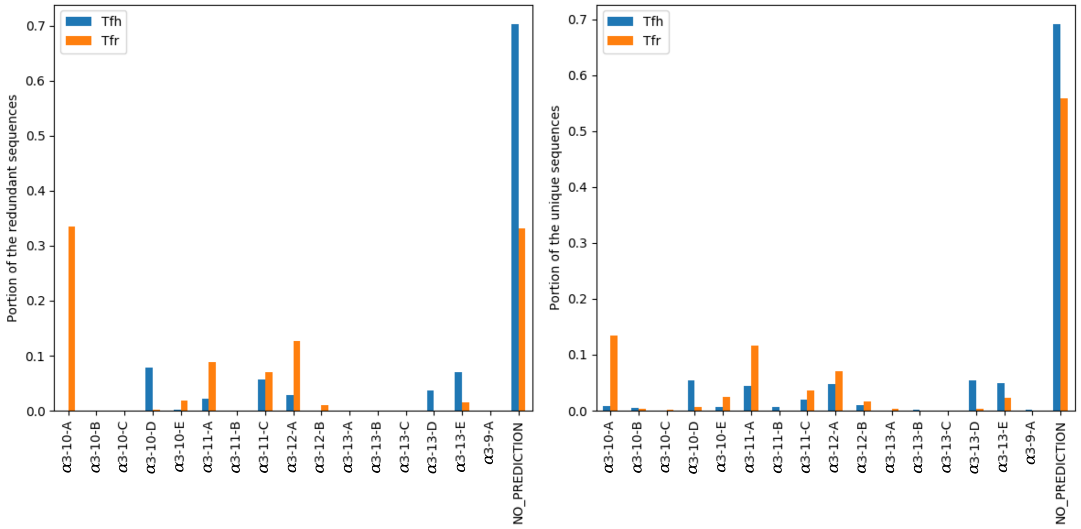


Figure S8: CDR$\alpha$3 canonical form prediction from the TCR sequences in Tfh and Tfr cells from (32).

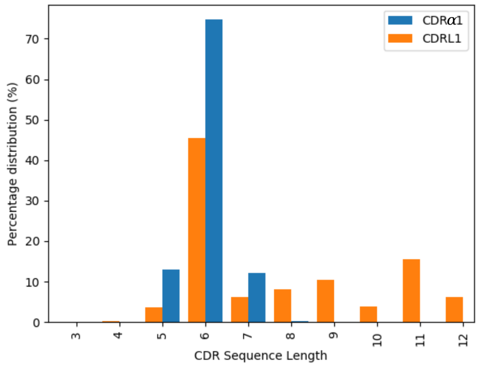


Figure S9: Length distributions of CDR$\alpha$1 (blue) and CDRL1 (orange) loops. The length distributions overlap between CDR$\alpha$1 and CDRL1.

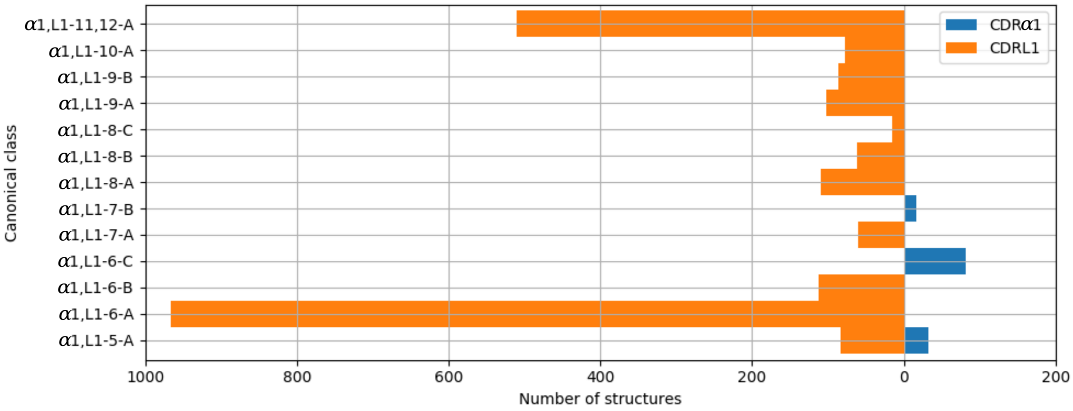


Figure S10: Clusters of CDR$\alpha$1 (blue) and CDRL1 (orange) loops. Orange bars indicate the number of CDRL1 structures, and blue bars the number of CDR$\alpha$1 structures. All classes, apart from $\alpha$1,L1-5-A, have structures from only one CDR type, *i.e.* CDR$\alpha$1 or CDRL1.

| Canonical class | Al-Lazikani | Klausen | Number of unique sequences (Number of structures) |
|---|---|---|---|
| $\beta$1-5-A | $\beta$1-1/$\beta$1-2 | $\beta$1-1/$\beta$1-2 | 30 (344) |
| $\beta$1-6-A* | $\beta$1-3 | | 1 (5) |
| (unclustered) | | $\beta$1-3/$\beta$1-4 | |
| $\beta$1-unclustered | | | 3 (8) |
| $\beta$2-6-A | $\beta$2-1 | $\beta$2-2 | 18 (238) |
| $\beta$2-6-B | | $\beta$2-3/$\beta$2-4/$\beta$2-5 | 16 (102) |
| $\beta$2-6-C* | $\beta$2-3 | $\beta$2-6 | 1 (6) |
| (unclustered) | $\beta$2-2 | $\beta$2-1/$\beta$2-7 | |
| $\beta$2-unclustered | | | 6 (14) |

Table S2: Summary of CDR$\beta$ classes. Clusters are labelled and mapped to previous canonical classes (see Table S1 and Materials and Methods). If there is at least one structure that is present in both our canonical class and those from the literature, we consider these classes to be analogous. Pseudo-classes are indicated with *.

Table S3: Number of sequences with multiple conformations.

| | TCR and antibody (antibody subset) | | | | | |
|---|---|---|---|---|---|---|
| | L1 | L2 | L3 | H1 | H2 | Total |
| # Unique sequences | 580 | 198 | 855 | 793 | 885 | 3311 |
| # Unique sequences with multiple structures | 362 | 151 | 527 | 503 | 546 | 2089 |
| # Unique sequences with structures in multiple classes | 25 | 12 | 17 | 42 | 70 | 166 |
| | TCR and antibody (TCR subset) | | | | | |
| | $\alpha$1 | $\alpha$2 | $\alpha$3 | $\beta$1 | $\beta$2 | Total |
| # Unique sequences | 45 | 44 | 99 | 33 | 39 | 260 |
| # Unique sequences with multiple structures | 39 | 35 | 72 | 30 | 36 | 212 |
| # Unique sequences with structures in multiple classes | 11 | 6 | 9 | 3 | 12 | 41 |

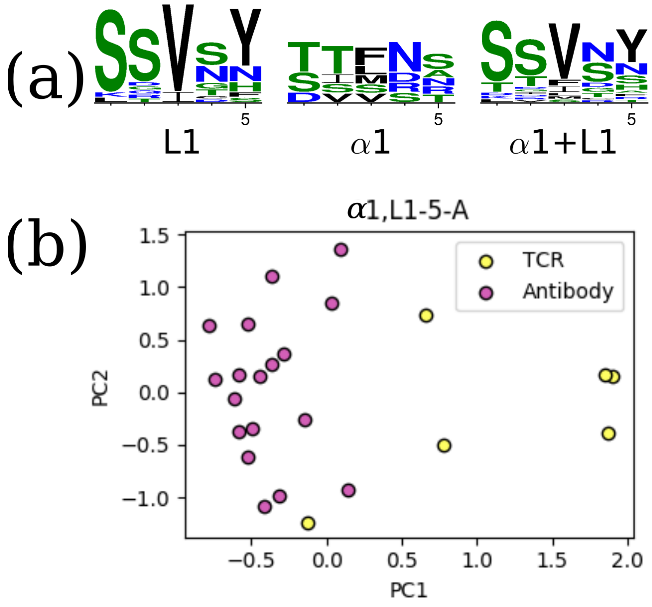


Figure S11: The unique TCR and antibody sequences in the $\alpha$1,L1-5-A class. (a) Sequence logos of CDR$\alpha$1 and CDRL1 loops in the $\alpha$1,L1-5-A class, using only the unique sequences. The sequence patterns are distinct between the two classes. (b) Principal component analysis (PCA) plot of the first two components in one-hot-encoded sequences, stratified by TCR and antibody CDRs (see Materials and Methods). TCR $\alpha$1 and antibody L1 can be separated by the first principal component. One TCR sequence (DSVNN) is close to the antibody set.

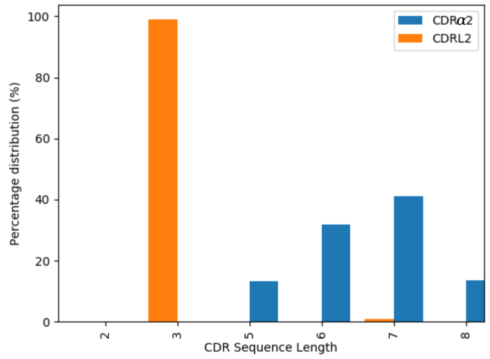


Figure S12: Length distribution of CDR$\alpha$2 (blue) and CDRL2 (orange) loops. The majority of the CDRL2 loops are three-residues long, while CDR$\alpha$2 adopts a range of sequence lengths.

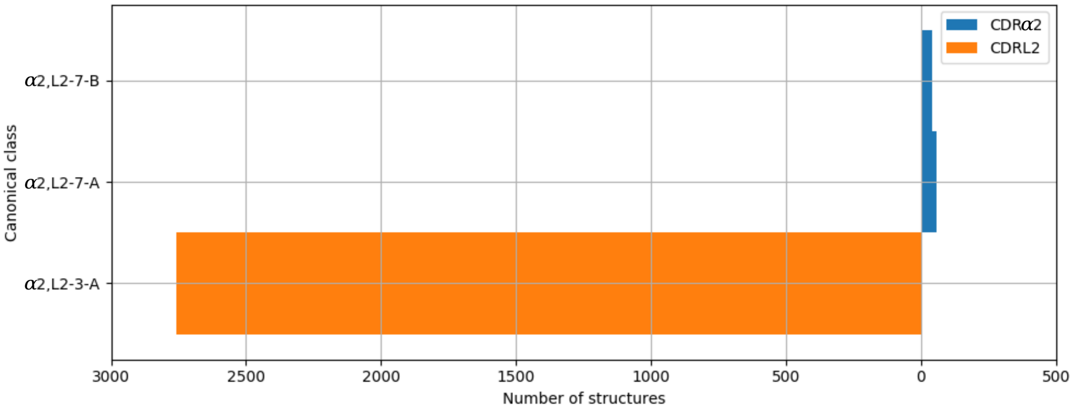


Figure S13: Clusters of CDR$\alpha$2 (blue) and CDRL2 (orange) loops. Orange bars indicate the number of CDRL2 structures, and blue bars the number of CDR$\alpha$2 structures. All clusters contain structures from only one CDR type, *i.e*. CDR$\alpha$2 or CDRL2.

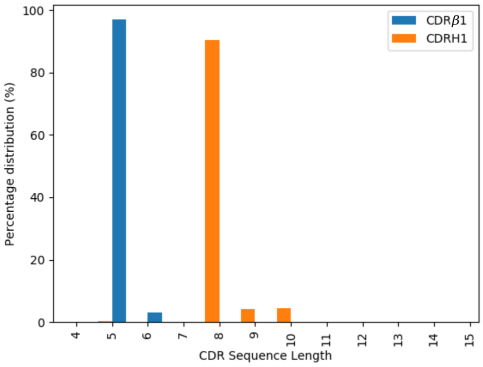


Figure S14: Length distributions of CDR$\beta$1 (blue) and CDRH1 (orange) loops. CDR$\beta$1 loops tend to be shorter than CDRH1 loops.

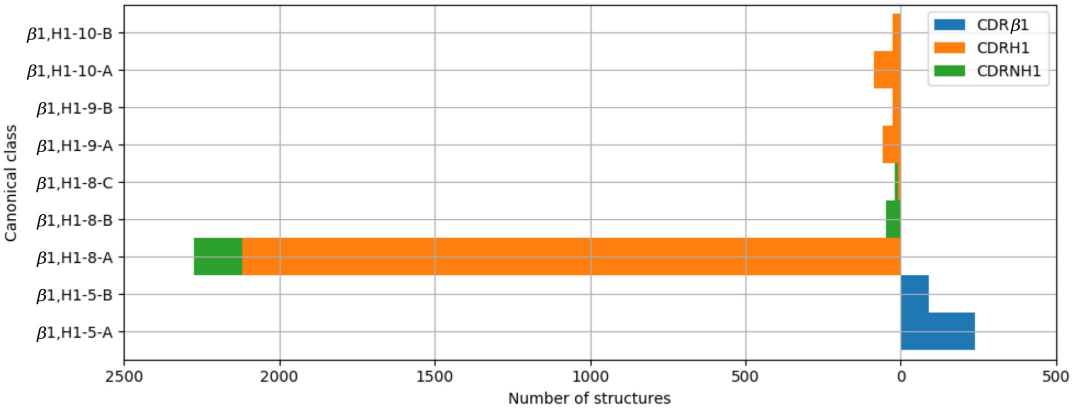


Figure S15: Clusters of CDR$\beta$1 and CDRH1 loops. Orange bars indicate the number of antibody CDRH1 structures, green bars nanobody CDRH1 structures, and blue bars the number of CDR. All classes have structures from only one CDR type, *i.e*. CDR$\beta$1 or CDRH1. Some nanobody CDRH1 loops are structurally closer to antibody CDRH1 loops than TCR CDR$\beta$1 loops.

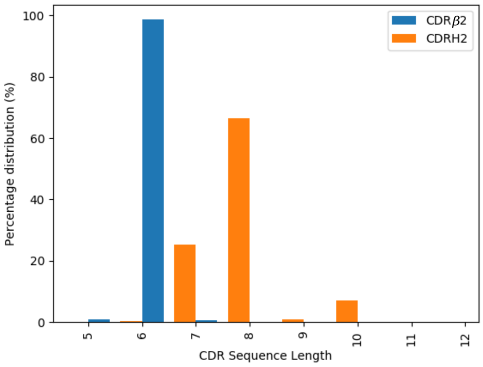


Figure S16: Length distributions of CDR$\beta$2 (blue) and CDRH2 (orange) loops. The majority of the CDR$\beta$2 loops are slightly shorter than CDRH2 loops.

.

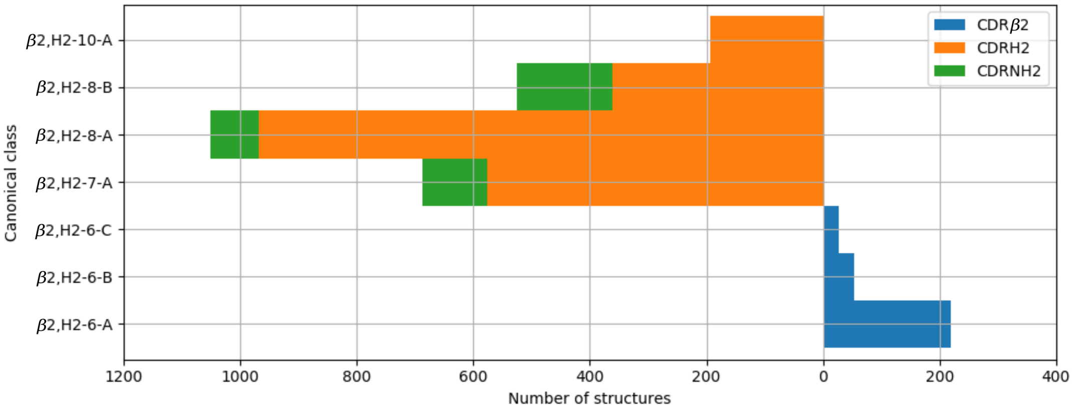


Figure S17: Clusters of CDR$\beta$2 and CDRH2 loops. Orange bars indicate the number of antibody CDRH2 structures, green bars nanobody CDRH2 structures, and blue bars the number of CDR$\beta$2 structures. All classes have structures from only one CDR type, *i.e*. CDR$\beta$2 or CDRH2. Antibody and nanobody CDRH2 loops are structurally more similar than TCR CDR$\beta$2 loops.

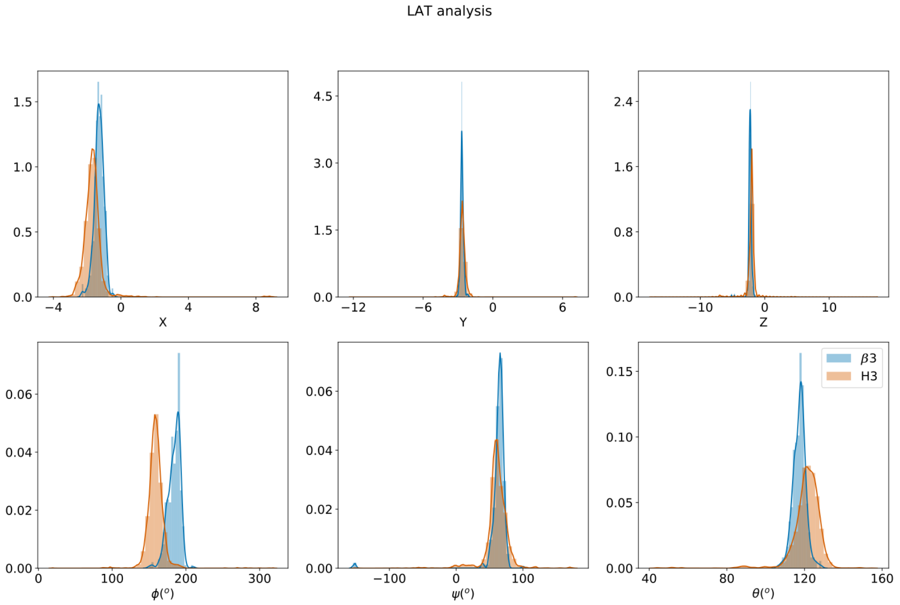


Figure S18: Loop anchor transformation (LAT) analysis of CDR$\beta$3 and CDRH3 structures. The Euler transformation was calculated on IMGT positions 105 and 117 (see Materials and Methods). X, Y, Z, $\phi$, $\psi$ and $\theta$ are the six degrees of freedom in the Euler transformation.

.

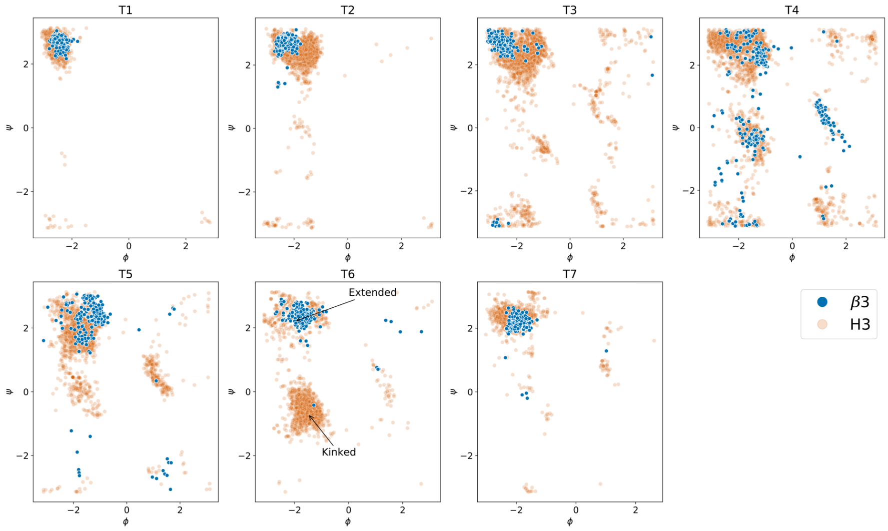


Figure S19: The $\phi/\psi$ plot of the first three (T1-T3, corresponding to IMGT positions 105-107) and last four (T4-T7, corresponding to IMGT positions 114-117) loop residues, in CDR$\beta$3 and CDRH3. Residue T6 (IMGT position 116) populate different regions of the plot depending on whether the torso is extended or kinked (as labelled). See Materials and Methods and (19) for details.

.

Table S4: TCR CDRs with multiple conformations that fall into different structural clusters (including canonical forms and pseudo-classes). 'B' and 'U' refer to the number of bound and unbound structures of the sequence within that cluster.

| CDR | Sequence | Cluster 1 | B | U | Cluster 2 | B | U | Cluster 3 | B | U |
|---|---|---|---|---|---|---|---|---|---|---|
| $\alpha$1 | ATGYPS | UC | 4 | - | $\alpha$1,L1-6-H* | - | 1 | - | - | - |
| | DRGSQS | UC | - | 2 | $\alpha$1,L1-6-C | 16 | 5 | - | - | - |
| | DRNFQY | UC | - | 1 | $\alpha$1,L1-6-C | - | 3 | - | - | - |
| | DSAIYN | UC | 1 | - | $\alpha$1,L1-6-J* | 13 | 10 | - | - | - |
| | DSSSTY | UC | 1 | - | $\alpha$1,L1-6-C | 1 | - | - | - | - |
| | DSVNN | UC | 1 | - | $\alpha$1,L1-5-A | 3 | 4 | $\alpha$1,L1-5-B* | 5 | 2 |
| | YSATPY | UC | - | 1 | $\alpha$1,L1-6-D* | 4 | - | $\alpha$1,L1-6-E* | 4 | - |
| | VSGLRG | UC | 3 | - | $\alpha$1,L1-6-G* | 6 | - | - | - | - |
| | TSGFNG | UC | 6 | 4 | $\alpha$1,L1-6-G* | 51 | - | - | - | - |
| | TISGTDY | UC | - | 1 | $\alpha$1,L1-7-E* | 12 | - | - | - | - |
| | SSVPPY | $\alpha$1,L1-6-D* | 2 | 5 | $\alpha$1,L1-6-E* | - | 13 | - | - | - |
| $\alpha$2 | QEAYKQQN | UC | 1 | - | $\alpha$2,L2-8-A* | 3 | - | - | - | - |
| | ATKADDK | UC | 4 | - | $\alpha$2,L2-7-B | 2 | 1 | - | - | - |
| | SSTDNKR | UC | - | 2 | $\alpha$2,L2-7-B | - | 2 | - | - | - |
| | YYSGDPVV | UC | 1 | - | $\alpha$2,L2-8-B* | 7 | 1 | - | - | - |
| | GLTSN | UC | 3 | 1 | $\alpha$2,L2-5-A* | 6 | - | - | - | - |
| | GLKNN | UC | - | 2 | $\alpha$2,L2-5-A* | 4 | 2 | - | - | - |
| A3 | AMRGDSSYKLI | UC | - | 1 | $\alpha$3,L3-11-B* | 15 | 1 | - | - | - |
| | AGAGSQGNLI | UC | 1 | 1 | $\alpha$3,L3-10-H* | 7 | 1 | - | - | - |
| | AVNFGGGKLI | UC | - | 2 | $\alpha$3,L3-10-J* | 5 | - | - | - | - |
| | GTYNQGGKLI | UC | 2 | - | $\alpha$3,L3-10-A | 1 | - | - | - | - |
| | AYGEDDKII | UC | 3 | - | $\alpha$3,L3-9-D* | 2 | - | - | - | - |
| | AVTTDSWGKLQ | UC | - | 2 | $\alpha$3,L3-11-J* | 9 | - | - | - | - |
| | AVSESPFGNEKLT | UC | - | 1 | $\alpha$3,L3-13-H* | 2 | 3 | - | - | - |
| | AVRPTSGGSYIPT | UC | - | 1 | $\alpha$3,L3-13-C* | 5 | - | - | - | - |
| | AVRPLLDGTYIPT | UC | - | 1 | $\alpha$3,L3-13-C* | 3 | - | - | - | - |
| $\beta$1 | SEHNR | $\beta$1,H1-5-B | 1 | 5 | $\beta$1,H1-5-D* | 2 | 1 | - | - | - |
| | MGHDK | UC | 1 | - | $\beta$1,H1-5-A | 7 | 6 | - | - | - |
| | ENHRY | UC | 2 | - | $\beta$1,H1-5-A | 4 | 1 | - | - | - |
| $\beta$2 | FQNEAQ | UC | 1 | - | $\beta$2,H2-6-C | 8 | 9 | - | - | - |
| | FNNNVP | UC | 1 | - | $\beta$2,H2-6-B | 15 | 2 | - | - | - |
| | FQNQEV | UC | - | 1 | $\beta$2,H2-6-D* | 4 | 3 | - | - | - |
| | FYNGKV | UC | - | 1 | $\beta$2,H2-6-B | - | 3 | - | - | - |
| | SASEGT | UC | 2 | - | $\beta$2,H2-6-A | 47 | - | - | - | - |
| | SFDVKD | $\beta$2,H2-6-A | - | 6 | $\beta$2,H2-6-F* | - | 6 | - | - | - |
| | SVGEGT | UC | - | 2 | $\beta$2,H2-6-A | 3 | 3 | - | - | - |
| | SYGAGN | UC | 2 | - | $\beta$2,H2-6-A | 2 | - | - | - | - |
| | SYGAGS | UC | 1 | - | $\beta$2,H2-6-A | 34 | 10 | - | - | - |
| | SYGVNS | UC | 3 | 4 | $\beta$2,H2-6-A | 5 | 2 | - | - | - |
| | FQGTGA | UC | 1 | - | $\beta$2,H2-6-C | 2 | - | - | - | - |
| | YYNGEE | UC | 1 | - | $\beta$2,H2-6-B | 14 | - | - | - | - |

Table S5: Antibody CDRs with multiple conformations that fall into different structural clusters (including canonical forms and pseudo-classes). 'B' and 'U' refer to the number of bound and unbound structures of the sequence within that cluster.

| CDR | Sequence | Cluster 1 | B | U | Cluster 2 | B | U | Cluster 3 | B | U |
|---|---|---|---|---|---|---|---|---|---|---|
| L1 | QDISNY | UC | - | 1 | α1,L1-6-A | 30 | 29 | - | - | - |
| | QDINSY | UC | 2 | - | α1,L1-6-A | 5 | 2 | - | - | - |
| | KSVSTSGYSY | UC | 1 | - | α1,L1-10-A | 1 | - | - | - | - |
| | KSVSTSGYNY | UC | 1 | - | α1,L1-10-A | 1 | - | - | - | - |
| | ESVDYYGKSF | UC | - | 2 | α1,L1-10-A | 2 | - | - | - | - |
| | ENVVTY | UC | - | 1 | α1,L1-6-A | 1 | 3 | - | - | - |
| | QSVSSSY | α1,L1-7-A | 6 | 2 | α1,L1-7-C* | - | 3 | - | - | - |
| | QNLLDSSFDTNT | UC | - | 1 | α1,L1-11,12-A | - | 1 | - | - | - |
| | QSVSSY | UC | - | 3 | α1,L1-6-A | 11 | 12 | - | - | - |
| | QSVTNY | UC | 1 | 1 | α1,L1-6-I* | 2 | - | - | - | - |
| | QSVTSSQ | UC | - | 1 | α1,L1-7-A | - | 1 | - | - | - |
| | SRVGY | UC | - | 1 | α1,L1-5-A | - | 3 | - | - | - |
| | SSDVGGYNY | UC | 22 | 4 | α1,L1-9-A | 15 | 13 | - | - | - |
| | SSNIGENS | UC | - | 1 | α1,L1-8-A | - | 5 | - | - | - |
| | QGIRND | UC | 1 | - | α1,L1-6-A | 2 | - | - | - | - |
| | SSVSSSY | UC | 2 | - | α1,L1-7-A | 4 | 4 | - | - | - |
| | SSVSY | UC | 2 | - | α1,L1-5-A | 23 | 16 | - | - | - |
| | SSVTY | UC | - | 1 | α1,L1-5-A | - | 7 | - | - | - |
| | TGAVTSGHY | UC | 1 | - | α1,L1-9-B | 7 | - | - | - | - |
| | SSNVGGYNY | UC | - | 1 | α1,L1-9-A | - | 1 | - | - | - |
| | QSIVHSNGNTY | UC | - | 1 | α1,L1-11,12-A | 30 | 17 | - | - | - |
| | QSLSSKY | UC | - | 1 | α1,L1-7-A | - | 1 | - | - | - |
| | QSLVHSNGNTY | UC | 1 | 1 | α1,L1-11,12-A | 26 | 16 | - | - | - |
| | QSVDYDGDSY | UC | - | 1 | α1,L1-10-A | 1 | 3 | - | - | - |
| | QSVSSA | UC | 11 | - | α1,L1-6-A | 22 | - | α1,L1-6-I* | 3 | - |

Continued on next page

Table S5 – continued from previous page

| CDR | Sequence | Cluster 1 | B | U | Cluster 2 | B | U | Cluster 3 | B | U |
|---|---|---|---|---|---|---|---|---|---|---|
| L2 | LTS | UC | - | 1 | $\alpha$2,L2-3-A | 5 | - | - | - | - |
| | GNN | UC | 4 | - | $\alpha$2,L2-3-A | 12 | 2 | - | - | - |
| | EVN | UC | - | 1 | $\alpha$2,L2-3-A | 35 | 25 | - | - | - |
| | GAS | UC | 1 | - | $\alpha$2,L2-3-A | 101 | 42 | - | - | - |
| | FTS | $\alpha$2,L2-3-A | 5 | 1 | $\alpha$2,L2-3-D* | - | 1 | - | - | - |
| | GKN | $\alpha$2,L2-3-E* | 6 | 1 | $\alpha$2,L2-3-B* | 2 | 5 | - | - | - |
| | DVN | UC | 2 | - | $\alpha$2,L2-3-A | 2 | 3 | - | - | - |
| | YTS | UC | - | 1 | $\alpha$2,L2-3-A | 64 | 37 | $\alpha$2,L2-3-D* | 1 | 3 |
| | DST | UC | 1 | - | $\alpha$2,L2-3-A | - | 1 | - | - | - |
| | SDS | $\alpha$2,L2-3-A | 1 | - | $\alpha$2,L2-3-B* | 1 | - | - | - | - |
| | DAS | UC | - | 1 | $\alpha$2,L2-3-A | 86 | 78 | - | - | - |
| | GDN | $\alpha$2,L2-3-A | 1 | - | A2,L2-3-B* | 1 | - | - | - | - |
| L3 | ALWYSNHLV | UC | 1 | 1 | $\alpha$3,L3-9-B | 13 | 8 | - | - | - |
| | YSTDSSGNHRV | UC | 1 | - | $\alpha$3,L3-11-A | - | 2 | - | - | - |
| | SSYGGDNNLF | UC | - | 1 | $\alpha$3,L3-10-A | - | 2 | - | - | - |
| | SSYEGSDNFV | UC | 17 | 4 | $\alpha$3,L3-10-C | 16 | 5 | - | - | - |
| | SSRDKSGSRLSV | UC | - | 1 | $\alpha$3,L3-12-C* | 5 | - | - | - | - |
| | QVYGASSYT | UC | 4 | - | $\alpha$3,L3-9-D* | - | 6 | - | - | - |
| | QQWSSHIFT | UC | - | 1 | $\alpha$3,L3-9-B | - | 1 | $\alpha$3,L3-9-C | - | 1 |
| | QQWNYPFT | UC | - | 1 | $\alpha$3,L3-8-D* | 2 | 1 | - | - | - |
| | QAWDASTGV | UC | 1 | - | $\alpha$3,L3-9-G* | 7 | - | - | - | - |
| | QQHYTTPPT | $\alpha$3,L3-9-A | 14 | 18 | $\alpha$3,L3-9-F* | - | 6 | - | - | - |
| | QAWDSSTVV | UC | - | 1 | $\alpha$3,L3-9-B | 1 | 2 | - | - | - |
| | HQYLSSWT | UC | - | 1 | $\alpha$3,L3-8-A | - | 2 | - | - | - |
| | GVGDTIKEQFVYV | UC | 2 | 3 | $\alpha$3,L3-13-F* | 5 | - | - | - | - |
| | GTWDSSLNPV | UC | - | 1 | $\alpha$3,L3-10-A | 1 | 1 | - | - | - |
| | AAWDDSLDVAV | UC | - | 2 | $\alpha$3,L3-10,11-A | - | 4 | - | - | - |
| | QQSSSSLIT | $\alpha$3,L3-9-C | 12 | - | $\alpha$3,L3-9-A | 1 | - | - | - | - |
| | FQGSHVPRT | UC | 1 | 1 | $\alpha$3,L3-9-A | 2 | 6 | - | - | - |
| H1 | RRSSRSWA | UC | - | 5 | $\beta$1,H1-8-D* | - | 2 | $\beta$1,H1-8-T* | 6 | - |
| | GRTFSSYG | UC | 1 | - | $\beta$1,H1-8-C | 3 | - | - | - | - |
| | GRTFSSYA | UC | - | 1 | $\beta$1,H1-8-A | 1 | - | $\beta$1,H1-8-C | 1 | - |
| | GHTYSTYC | UC | 1 | 2 | $\beta$1,H1-8-F* | 2 | - | - | - | - |
| | GGTFSSYA | UC | 3 | 4 | $\beta$1,H1-8-A | 2 | 2 | - | - | - |
| | GGSISTYY | UC | 1 | - | $\beta$1,H1-8-A | 1 | - | - | - | - |
| | GGSISSSSYY | UC | - | 1 | $\beta$1,H1-10-B | - | 1 | - | - | - |
| | GGSFSTYA | UC | - | 3 | $\beta$1,H1-8-A | 28 | 2 | - | - | - |
| | GRTFSSYV | $\beta$1,H1-8-A | - | 1 | $\beta$1,H1-8-C | 7 | - | - | - | - |
| | GGSFSSYY | UC | 1 | - | $\beta$1,H1-8-A | 2 | - | - | - | - |
| | GFTFSNYG | UC | 1 | - | $\beta$1,H1-8-A | 15 | 4 | - | - | - |
| | GFTFSDYD | UC | 3 | - | $\beta$1,H1-8-A | 1 | - | - | - | - |
| | GFSLSTSGIG | UC | 1 | - | $\beta$1,H1-10-A | 1 | - | - | - | - |
| | GFSLSDKA | UC | - | 2 | $\beta$1,H1-8-A | - | 1 | - | - | - |
| | GFSLRTSRVG | UC | 1 | 2 | $\beta$1,H1-10-C* | 1 | - | - | - | - |
| | GFSFNTNA | UC | 1 | 1 | $\beta$1,H1-8-A | 2 | 2 | - | - | - |
| | GFNIKDTY | UC | 1 | 1 | $\beta$1,H1-8-A | 37 | 36 | - | - | - |
| | GFTLDDYA | UC | 1 | - | $\beta$1,H1-8-A | 2 | - | - | - | - |
| | GSAVSDYA | UC | - | 1 | $\beta$1,H1-8-S* | - | 5 | - | - | - |
| | GSISGIVV | UC | - | 1 | $\beta$1,H1-8-B | 3 | - | - | - | - |
| | GSSFTGYN | UC | 1 | - | $\beta$1,H1-8-A | 2 | - | - | - | - |

Continued on next page

Table S5 – continued from previous page

| CDR | Sequence | Cluster 1 | B | U | Cluster 2 | B | U | Cluster 3 | B | U |
|---|---|---|---|---|---|---|---|---|---|---|
| | GYTFTSYW | UC | 4 | - | $\beta$1,H1-8-A | 34 | 30 | - | - | - |
| | GYTFTSNW | $\beta$1,H1-8-A | 1 | 6 | $\beta$1,H1-8-C | - | 1 | - | - | - |
| | GYTFTSHW | UC | - | 1 | $\beta$1,H1-8-A | - | 1 | - | - | - |
| | GYTFTNYY | UC | 1 | 1 | $\beta$1,H1-8-A | 4 | 1 | - | - | - |
| | GYTFTNYG | UC | 1 | 1 | $\beta$1,H1-8-A | 14 | 3 | - | - | - |
| | GYTFTEYF | UC | - | 1 | $\beta$1,H1-8-A | 3 | - | - | - | - |
| | GYTFSEYW | UC | - | 1 | $\beta$1,H1-8-A | 2 | - | - | - | - |
| | GYSLSTSGMG | UC | - | 1 | $\beta$1,H1-10-A | - | 1 | - | - | - |
| | GYSITTNYA | $\beta$1,H1-9-D* | 6 | - | $\beta$1,H1-9-B | 1 | 1 | - | - | - |
| | GYSITSNYA | $\beta$1,H1-9-B | 2 | - | $\beta$1,H1-9-A | - | 1 | - | - | - |
| | GYSITSGYS | $\beta$1,H1-9-A | 4 | - | $\beta$1,H1-9-C* | 3 | 2 | - | - | - |
| | GYSITSDYA | UC | 2 | - | $\beta$1,H1-9-A | 15 | 5 | - | - | - |
| | GYSITSDFA | $\beta$1,H1-9-B | 4 | 7 | $\beta$1,H1-9-A | 1 | 3 | - | - | - |
| | GYSITGGYS | UC | - | 2 | $\beta$1,H1-9-A | - | 6 | - | - | - |
| | GYAFSSSW | UC | - | 2 | $\beta$1,H1-8-A | 3 | 1 | - | - | - |
| | GVTFSNVA | UC | 1 | - | $\beta$1,H1-8-A | 1 | - | - | - | - |
| | GVRLSAYD | UC | - | 1 | $\beta$1,H1-8-A | - | 1 | - | - | - |
| | GFNFSSSS | UC | 1 | - | $\beta$1,H1-8-A | 2 | - | - | - | - |
| | GDSITSGY | UC | 1 | - | $\beta$1,H1-8-A | 17 | 2 | - | - | - |
| | GSISSITT | UC | 1 | - | $\beta$1,H1-8-B | - | 2 | - | - | - |
| | GDSITSAY | UC | 1 | - | B1,H1-8-A | 1 | - | - | - | - |
| H2 | ILPGSDST | UC | 1 | 2 | $\beta$2,H2-8-A | 1 | - | - | - | - |
| | IGPSGGIT | UC | - | 4 | $\beta$2,H2-8-B | - | 4 | - | - | - |
| | IDPSNGDT | UC | - | 1 | $\beta$2,H2-8-A | - | 1 | - | - | - |
| | IDPNSGGT | UC | 2 | - | $\beta$2,H2-8-A | 11 | 8 | - | - | - |
| | IDPNGGGT | UC | 1 | - | $\beta$2,H2-8-A | 6 | 1 | - | - | - |
| | IDPEQGNT | UC | - | 2 | $\beta$2,H2-8-A | 1 | 1 | - | - | - |
| | ISAGGDKT | UC | 2 | - | $\beta$2,H2-8-B | 2 | 2 | - | - | - |
| | IGSSGGQT | $\beta$2,H2-8-B | 1 | - | $\beta$2,H2-8-C* | - | 1 | - | - | - |
| | ISASGGST | $\beta$2,H2-8-B | - | 4 | $\beta$2,H2-8-G* | 2 | - | - | - | - |
| | ISGSGGNT | UC | 1 | - | $\beta$2,H2-8-B | - | 1 | - | - | - |
| | FNPSNGRT | UC | - | 1 | $\beta$2,H2-8-A | - | 1 | - | - | - |
| | ISGSGGST | UC | 1 | - | $\beta$2,H2-8-A | - | 2 | $\beta$2,H2-8-B | 25 | 21 |
| | ASNGGII | UC | - | 1 | $\beta$2,H2-7-A | - | 1 | - | - | - |
| | ISNLDGST | UC | 2 | 1 | $\beta$2,H2-8-A | - | 1 | - | - | - |
| | ISPGNGDI | UC | - | 1 | $\beta$2,H2-8-A | 2 | - | - | - | - |
| | ISPYSGVT | UC | 1 | - | $\beta$2,H2-8-A | 1 | 1 | - | - | - |
| | ISGGGRNI | $\beta$2,H2-8-B | - | 1 | $\beta$2,H2-8-L* | 4 | - | - | - | - |
| | IGTSGNI | UC | 1 | 1 | $\beta$2,H2-7-A | 4 | 1 | - | - | - |
| | IHYRGTT | UC | - | 1 | $\beta$2,H2-7-A | 1 | 1 | - | - | - |
| | IIPIFGTA | UC | 3 | 2 | $\beta$2,H2-8-A | 4 | 6 | - | - | - |
| | IRSKSINSAT | UC | 1 | - | $\beta$2,H2-10-A | - | 3 | - | - | - |
| | IRSGGGRT | UC | 3 | - | $\beta$2,H2-8-B | 1 | - | - | - | - |
| | IRNKPYNYET | UC | 1 | - | $\beta$2,H2-10-A | 2 | - | - | - | - |
| | INTHSGVP | UC | 1 | - | $\beta$2,H2-8-A | 1 | - | - | - | - |
| | INSNGGST | UC | 1 | - | $\beta$2,H2-8-B | 6 | 1 | - | - | - |
| | INPGSDYT | UC | 2 | - | $\beta$2,H2-8-A | 2 | 1 | - | - | - |
| | INPGNGYT | UC | - | 2 | $\beta$2,H2-8-A | 2 | 1 | - | - | - |
| | INLNGGRT | UC | - | 1 | $\beta$2,H2-8-A | - | 1 | - | - | - |
| | INIYTGEP | UC | 1 | - | $\beta$2,H2-8-A | 2 | 1 | - | - | - |

Continued on next page

Table S5 – continued from previous page

| CDR | Sequence | Cluster 1 | B | U | Cluster 2 | B | U | Cluster 3 | B | U |
|---|---|---|---|---|---|---|---|---|---|---|
| | INIGATYA | UC | 1 | - | $\beta$2,H2-8-B | 1 | - | - | - | - |
| | ILPGSGST | UC | 3 | - | $\beta$2,H2-8-A | 4 | 5 | - | - | - |
| | ILPGSGRT | UC | 2 | - | $\beta$2,H2-8-A | 1 | 1 | - | - | - |
| | ILPGGGSN | UC | - | 2 | $\beta$2,H2-8-A | - | 2 | - | - | - |
| | IKSKTDGGTT | UC | - | 1 | $\beta$2,H2-10-A | - | 5 | - | - | - |
| | IIPILGIA | UC | 1 | - | $\beta$2,H2-8-A | 1 | - | - | - | - |
| | ISRSGSVT | UC | 1 | - | $\beta$2,H2-8-B | 2 | 1 | - | - | - |
| | IRSSVIT | UC | 1 | - | $\beta$2,H2-7-B* | 6 | 1 | - | - | - |
| | ISSGGGNT | UC | - | 2 | $\beta$2,H2-8-B | 1 | - | - | - | - |
| | ISSPGTI | UC | - | 3 | $\beta$2,H2-7-A | - | 1 | - | - | - |
| | VIPLLTIT | UC | 4 | - | $\beta$2,H2-8-A | 25 | 6 | - | - | - |
| | TNPRNGGT | UC | 1 | - | $\beta$2,H2-8-A | 4 | 2 | - | - | - |
| | TIPLFGKT | UC | - | 1 | $\beta$2,H2-8-A | 1 | 1 | - | - | - |
| | NNPGNGYI | UC | - | 1 | $\beta$2,H2-8-A | - | 1 | - | - | - |
| | MSHEGDKT | UC | - | 1 | $\beta$2,H2-8-B | - | 1 | - | - | - |
| | MDSGGGGT | UC | 9 | 1 | $\beta$2,H2-8-B | - | 1 | - | - | - |
| | LTTTGTA | $\beta$2,H2-7-A | 3 | 1 | $\beta$2,H2-7-C* | 1 | 4 | - | - | - |
| | IYWDDVE | UC | - | 1 | $\beta$2,H2-7-A | 1 | - | - | - | - |
| | IYWDDDK | UC | 1 | - | $\beta$2,H2-7-A | 8 | 7 | - | - | - |
| | IYSYYGST | UC | 2 | - | $\beta$2,H2-8-A | 2 | - | - | - | - |
| | IYSSYSYT | UC | 1 | - | $\beta$2,H2-8-A | 1 | - | - | - | - |
| | ISSGGGRT | UC | 1 | 1 | $\beta$2,H2-8-B | 1 | - | - | - | - |
| | IYPYYGST | $\beta$2,H2-8-A | 1 | - | $\beta$2,H2-8-D* | 3 | - | - | - | - |
| | IYPYSGST | $\beta$2,H2-8-A | 2 | - | $\beta$2,H2-8-D* | 1 | - | - | - | - |
| | IYRSGSRM | UC | - | 1 | $\beta$2,H2-8-B | - | 1 | - | - | - |
| | IYPTNGYT | UC | 1 | - | $\beta$2,H2-8-A | 16 | 28 | - | - | - |
| | ISWSGGST | UC | - | 1 | $\beta$2,H2-8-B | 1 | 1 | - | - | - |
| | ITGPGEGWSV | UC | 1 | - | $\beta$2,H2-10-A | 9 | - | - | - | - |
| | ITPAGGYT | UC | 3 | - | $\beta$2,H2-8-J* | - | 18 | - | - | - |
| | VYDSGDT | UC | - | 3 | $\beta$2,H2-7-A | - | 1 | - | - | - |
| | IWAGGST | UC | 2 | - | $\beta$2,H2-7-A | - | 1 | - | - | - |
| | IWGDGIT | UC | - | 1 | $\beta$2,H2-7-A | 1 | - | - | - | - |
| | ITYSGTT | UC | 1 | 1 | $\beta$2,H2-7-A | 1 | - | - | - | - |
| | IWSGGST | UC | - | 1 | $\beta$2,H2-7-A | 7 | 10 | - | - | - |
| | IWWDDDN | UC | 1 | - | $\beta$2,H2-7-A | 1 | - | - | - | - |
| | IWYSGSNT | UC | - | 2 | $\beta$2,H2-8-C* | - | 4 | $\beta$2,H2-8-K* | - | 9 |
| | IYPGNGDT | UC | 1 | - | $\beta$2,H2-8-A | 4 | 11 | - | - | - |
| | IYPGNVHA | UC | - | 1 | $\beta$2,H2-8-A | - | 2 | - | - | - |
| | IYPGSGGT | UC | - | 1 | $\beta$2,H2-8-A | - | 2 | - | - | - |
| | IWPSGGNT | UC | 1 | - | $\beta$2,H2-8-B | 1 | 1 | - | - | - |
| | IRWNGGST | UC | 1 | - | $\beta$2,H2-8-B | 2 | - | - | - | - |

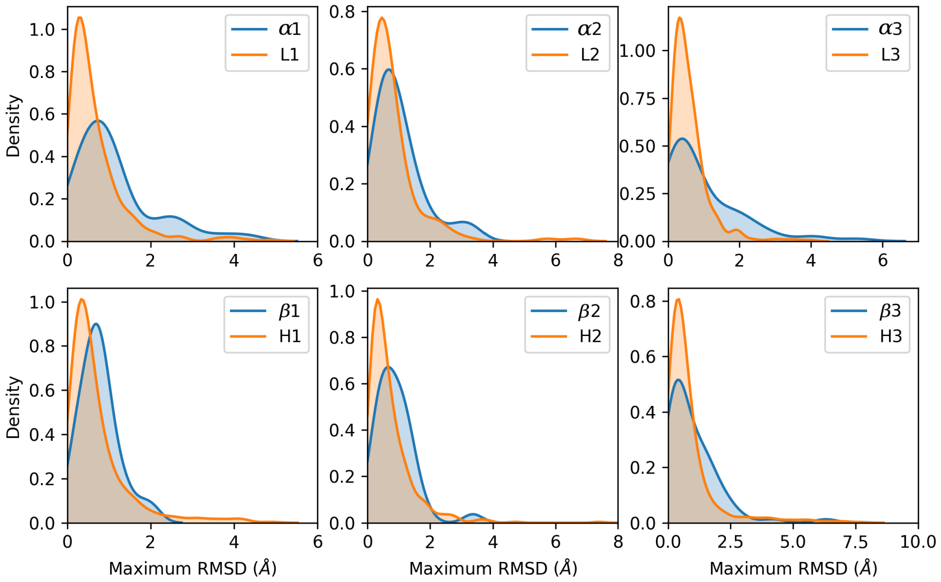


Figure S20: The density of the maximum root-mean square deviation (RMSD) between loop structures with identical CDR sequence. TCR CDRs (blue) show a shift to the right compared to antibody CDRs (orange), suggesting that structural flexibility of TCR CDRs is greater than that of antibodies.

.
